# memento: Generalized differential expression analysis of single-cell RNA-seq with method of moments estimation and efficient resampling

**DOI:** 10.1101/2022.11.09.515836

**Authors:** Min Cheol Kim, Rachel Gate, David S. Lee, Andrew Lu, Erin Gordon, Eric Shifrut, Alexander Marson, Vasilis Ntranos, Chun Jimmie Ye

**Affiliations:** Medical Scientist Training Program, University of California, San Francisco, San Francisco, California, USA; UC Berkeley-UCSF Graduate Program in Bioengineering, San Francisco, CA, USA; Institute for Human Genetics, University of California, San Francisco, San Francisco, California, USA; Division of Pulmonary and Critical Care, University of California, San Francisco, San Francisco, California, USA; Diabetes Center, University of California, San Francisco, San Francisco, CA, USA; Department of Microbiology and Immunology, University of California San Francisco, San Francisco, CA, USA; Institute of Computational Health Sciences, University of California, San Francisco, San Francisco, CA, USA; Department of Bioengineering and Therapeutic Sciences, University of California, San Francisco, San Francisco, CA, USA; Department of Epidemiology and Biostatistics, University of California, San Francisco, San Francisco, CA, USA; Chan Zuckerberg Biohub, San Francisco, CA, USA; Parker Institute for Cancer Immunotherapy, San Francisco, CA, USA; Gladstone-UCSF Institute of Genomic Immunology, San Francisco, CA, USA; Division of Rheumatology, Department of Medicine, University of California, San Francisco, San Francisco, CA, USA

## Abstract

Differential expression analysis of scRNA-seq data is central for characterizing how experimental factors affect the distribution of gene expression. However, it remains challenging to distinguish biological and technical sources of cell-cell variability and to assess the statistical significance of quantitative comparisons between groups of cells. We introduce `memento` to address these limitations and enable accurate and efficient differential expression analysis of the mean, variability, and gene correlation from scRNA-seq. We used `memento` to analyze 70,000 tracheal epithelial cells to identify interferon response genes with distinct variability and correlation patterns, 160,000 T cells perturbed with CRISPR-Cas9 to reconstruct gene-regulatory networks that control T cell activation, and 1.2 million PMBCs to map cell-type-specific *cis* expression quantitative trait loci (eQTLs). In all cases, `memento` identified more significant and reproducible differences in mean expression but also identified differences in variability and gene correlation that suggest distinct modes of transcriptional regulation imparted by cytokines, genetic perturbations, and natural genetic variation. These results demonstrate `memento` as a first-in-class method for the quantitative comparisons of scRNA-seq data scalable to millions of cells and thousands of samples.

## Introduction

Gene expression is determined by a cell’s genetic makeup and environmental exposure but can fluctuate because of extrinsic noise due to the specific state of a cell or intrinsic noise due to mRNA transcription and degradation [1, 2]. While genetics and environmental history account for much of expression variability in a population of cells, transcriptional noise has been shown to have profound effects on cellular response to perturbations and cellular development and differentiation [3, 2, 4]. Characterizing how deterministic and stochastic factors together shape the distribution of gene expression is central for understanding how transcriptional control is established, maintained, and may be broken. These insights, in turn, has the potential to identify mechanisms underlying phenomena where genotype-phenotype relationships are not completely explained, such as destabilization [3], incomplete penetrance [5], and variable expressivity [6].

The distribution of an individual gene’s expression over a population of cells has historically been parameterized by the mean, variance, and their derivatives such as the Fano factor and coefficient of variation [7]. The expression of constitutively expressed genes (e.g., housekeeping genes) that are transcribed and degraded at constant rates are expected to follow a Poisson distribution. However, foundational work in prokaryotes and yeast has observed that the expression for most genes in their genomes is over-dispersed, exhibiting higher than expected variability [8]. Furthermore, genes that participate in the same biological pathway have been demonstrated to be transcriptionally correlated [5]. These observations are consistent with a model of active regulation of multiple related genes each controlled by *cis* regulatory elements for the same transcription factors with “on” and “off” states [9]. Until recently, studying the distribution of gene expression, in particular the joint distribution of multiple genes, has been technologically challenging and has been mostly pursued in model organisms that can be genetically modified [10, 11].

Single-cell RNA-sequencing (scRNA-seq) has emerged as a systematic and efficient approach for profiling the transcriptomes of cells across experimental factors including extracellular stimuli [12], genetic perturbations [13, 14], and natural genetic variation [15, 16, 17, 18]. Analysis of scRNA-seq data can in theory determine how experimental factors and transcriptional noise together shape the distribution of gene expression across the domains of life. However, there remains a need for differential expression analysis methods that compare distributional parameters between groups of cells including the mean, variability, and gene correlation. To assess differences in mean expression, it is common practice to apply differential expression analysis methods for bulk RNA-seq to pseudobulk profiles generated by aggregating transcript counts for groups of cells defined by clustering. While pseudobulk approaches do not fully take advantage of single cells as repeated measures, they have been demonstrated to outperform methods that explicitly model the distribution of observed scRNA-seq data [19]. Very few methods exist for assessing differences in the variability of gene expression that measure transcriptional noise or correlation between pairs of genes that measure the coordinated expression of genes that may participate in the same regulatory network.

Generalized differential expression analysis of scRNA-seq data remains challenging due to two statistical limitations. First, decomposing the overall cell-to-cell variability from scRNA-seq data into transcriptional versus measurement noise remains difficult [20]. This is because small numbers of molecules are involved in the biochemical reactions of both gene transcription and the sampling process of scRNA-seq (**Fig. 1A**) [21]. Most existing methods implement highly parameterized models aimed to explain the higher than expected cell-cell variability in the *observed* sparse transcript counts but does not explicitly distinguish between biological versus technical sources of variability [22, 23, 24, 25, 26, 27]. Accurate estimates of biological variability is crucial for modeling joint distribution of multiple genes such as the correlation between pairs of genes [22]. Second, establishing whether a particular comparison of mean, variability, or gene correlation from scRNA-seq data is statistically significant remains a largely unsolved problem. Existing methods utilize asymptotic theory to establish statistical significance for the comparison of means which can result in p-value distributions that are either inflated or deflated. This is particularly problematic in settings where thousands of comparisons are made since poorly calibrated p-values violate assumptions for multiple testing correction. Further, most methods are not able to explicitly account for biological or technical replicates that sample multiplexed workflows can routinely generate with a growing number of individuals or conditions [28, 15, 29, 13, 30]. Indeed, recent studies have shown that scRNA-seq methods surprisingly underperform pseudobulk methods for testing mean differences likely due to limitations in both multiple testing correction and properly accounting for replicates [19].

**Figure 1:**
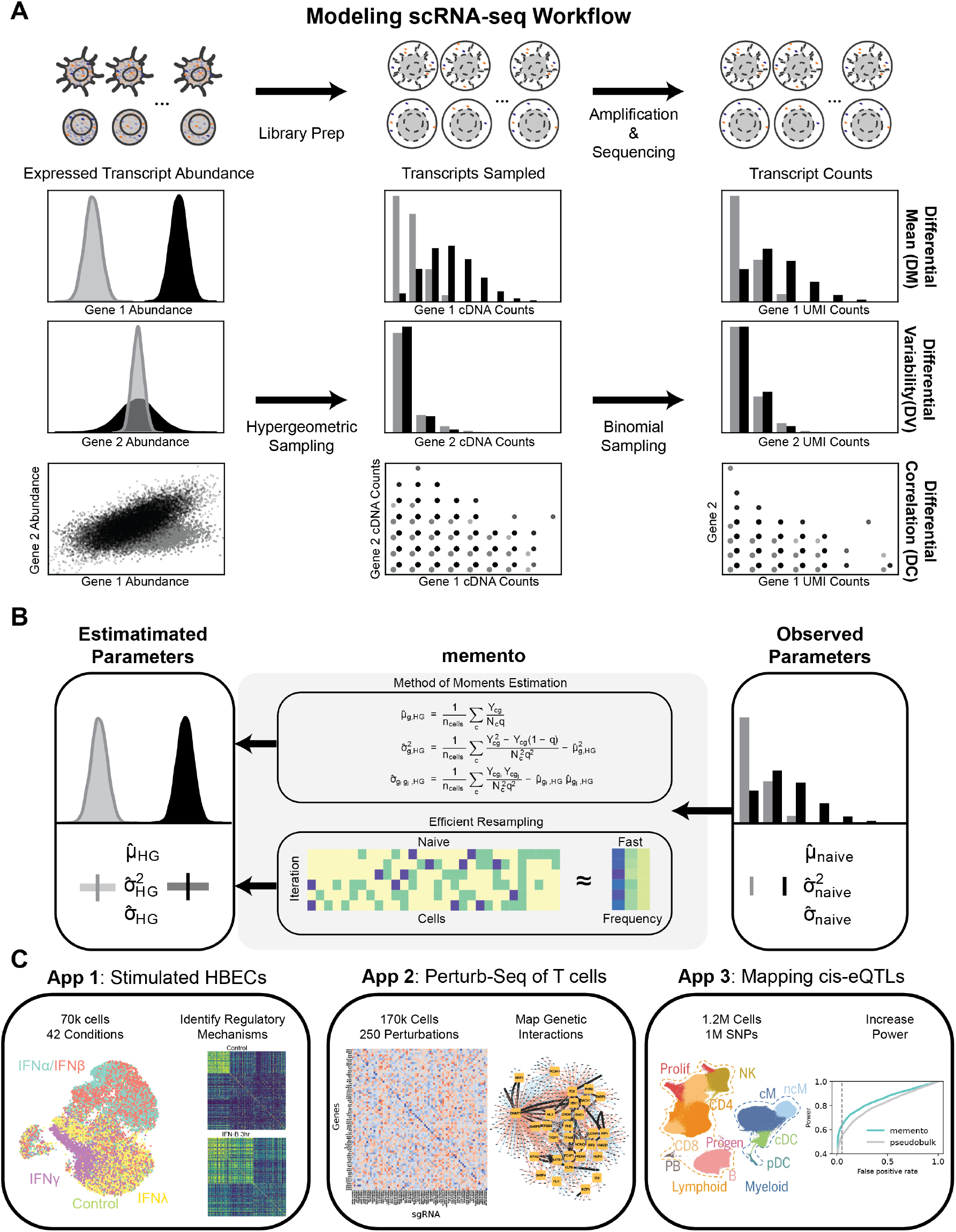
memento workflow for differential mean, variability, and gene correlation testing. **(A)** Experimental workflow for single cell RNA-sequencing samples RNA transcripts inside each cell during library preparation and sequencing. After scRNA-seq sampling, patterns of mean, variability, and correlation of gene expression in the observed transcript counts no longer resemble the actual distribution. **(B)** memento models scRNA-seq as a hypergeometric sampling process, estimates expression distribution parameters (mean, residual variance, and correlation) using method of moments estimators, implements efficient bootstrapping for estimating confidence intervals, and tests for differences in expression parameters between two groups of cells. **(C)** Three applications of memento to characterize the response of 70k human tracheal epithelial cells to extracellular cytokines, reconstruct the gene regulatory network from 170k human CD4+ T cells perturbed by CRISPR-Cas9, and map the genetic control of gene expression in 1.2M peripheral blood mononuclear cells from 162 SLE patients and 99 healthy controls.

To address these issues, we introduce `memento`, an end-to-end method that implements a hierarchical model for estimating the mean, residual variance, and gene correlation from scRNA-seq data and a statistical framework for hypothesis testing of differences in these parameters between groups of cells (**Fig. 1B**). `memento`models scRNA-seq using a novel multivariate hypergeometric sampling process while making no assumptions about the true distributional form of gene expression within cells. Importantly, by exploiting the sparsity of scRNA-seq data, `memento` implements an innovative bootstrapping strategy for efficient statistical comparisons of the estimated parameters between groups of cells that can also incorporate biological and technical replicates. Through simulations and analyses of real data, we demonstrate that `memento` produces accurate parameter estimates over a range of gene expression distributions and sampling efficiencies, computes well-calibrated test statistics suitable for multiple testing correction, and achieves sublinear runtimes scalable to the analysis of millions of cells. We demonstrate the broad applicability of `memento` in three applications to study how experimental factors affect the distribution of gene expression in human cells (**Fig. 1C**). First, we performed scRNA-seq on 70k tracheal epithelial cells stimulated with extracellular interferons and investigated how stimulation shapes the variability and correlation of response genes over time. Second, we performed Perturb-seq on 160k T cells and mapped gene regulatory networks that define broad T cell activation. Finally, we reanalyzed 1.2M cells collected from 250 individuals to identify genetic variants associated with mean, variability, and gene correlation in specific cell types. In all cases, `memento` identified more significant and reproducible differences in mean expression compared to existing methods but also identified differences in variability and gene correlation that revealed distinct modes of transcriptional regulation imparted by cytokines, genetic perturbations, and natural genetic variation. `memento` is implemented in python, is compatible with scanpy [31], and can be downloaded at https://github.com/yelabucsf/scrna-parameter-estimation.

## Results

### Novel statistical model of single-cell RNA-sequencing

It has long been observed that scRNA-seq produces sparse data displaying a high degree of cell-to-cell variability even in genetically identical cells exposed to the same environment (**Fig. 1A**). Decomposing this variability into biological versus measurement noise is critical to differential expression analysis of scRNA-seq data. Measurement noise present in scRNA-seq can be associated with inefficiencies in at least three molecular biology processes common to nearly all workflows: 1) only a fraction of the expressed transcripts is captured within compartments and undergoes reverse transcription (RT) to generate cDNA, 2) only a fraction of cDNA molecules is amplified during each round of polymerase chain reaction (PCR), and 3) only a fraction of the amplified cDNA is ultimately sequenced. The introduction of Unique Molecular Identifiers (UMIs) has largely obviated the need to model the noise introduced by PCR [32]. However, noise from imperfect transcript capture for RT and imperfect cDNA sampling during sequencing remain in the observed, attenuated distribution of counts.

We introduce a novel statistical model to describe the observed scRNA-seq counts as the result of hyper-geometric sampling of the expressed transcripts within a cell. The intuition for the hypergeometric model comes from the observation that the capturing of poly-adenylated mRNA for RT and sequencing of the resulting libraries sample molecules from each cell without replacement, and these sampling steps collectively create measurement noise in the final dataset. Key to our model is we allow for arbitrary distributions of gene expression within a cell prior to measurement. Formally, let 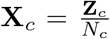 be a *m*-dimensional random variable representing the normalized transcript counts of *m* genes in cell *c*, where **Z**_*c*_ is a vector of the expressed transcript counts and *N*_*c*_ is the total transcript counts within a cell. We assume scRNA-seq to be a multivariate hypergeometric sampling process that produces the observed transcript counts **Y**_*c*_ from **X**_*c*_: **Y**_*c*_ *∼* MultiHG(*N*_*c*_**X**_*c*_, *N*_*c*_, *N*_*c*_*q*). In our formulation, *q* is the overall transcript sampling efficiency of scRNA-seq and can be related to measurement noise introduced during library preparation and sequencing (see **Methods**). We also empirically demonstrate that the two-step noise process of RT capture (hypergeometric) and sequencing (binomial) can be well represented with a single step of hypergeometric sampling with the overall *q* (**Fig. S1**).

### Estimating distributional parameters of gene expression from scRNA-seq

To the best of our knowledge, a hypergeometric sampling process has not been used to model scRNA-seq data for differential expression analysis. This may be partly because of challenges in deriving estimates of distribution parameters using maximum likelihood. Here, we derive method of moment (MoM) estimators for the first (mean), the second (variance), and the mixed (covariance) moment of **X**_*c*_ given **Y**_*c*_ assuming hypergeometric sampling (see **Methods** for derivation):

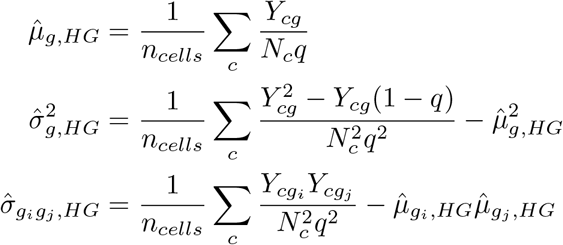

While the mean can be directly used to test for differential mean expression (DM), the variance need to be adjusted to account for the expected dependence between mean and variance in count based data in order to test for differential expression variability (DV) independent of DM [33, 34]. We compute the *residual variance* 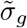 as a measure of expression variability *σ*_*g*_(**Methods**), defined as the component of variance that is not explained by the mean (**Methods**) and gene correlation as covariance terms (off diagonal elements) scaled by the variance terms (diagonal elements) from the variance-covariance matrix estimated above.

We performed extensive simulations to compare `memento` ’s hypergeometric estimators to the naive plug-in estimators (used by scHOT [35]), the empirical Bayes estimators using the Poisson approximation introduced by Zhang et al. [36], and estimates derived from BASiCS [27] (see **Methods** for forms of the naive and Poisson estimators). Across a range of reported *q*s from both low-efficiency (*q* < 0.2) droplet-based (e.g., 10X V1, V2 and V3) and high-efficiency (*q* > 0.3) plate-based (Smart-Seq3 [37]) scRNA-seq workflows, `memento` ’s hypergeometric estimator produced very accurate estimates of mean (Lin’s concordance correlation coefficient - *ρ*_*c*_ *>* 0.98), residual variance (*ρ*_*c*_ *>* 0.98), and gene correlation (*ρ*_*c*_ *>* 0.98) (**Fig. 2A**). While mean estimates were very similar for all estimators, `memento` produces stable residual variance and gene correlation estimates across *q*s, outperforming other estimators for both low- and high-efficiency workflows. These simulations assume sequencing is performed to saturation but as previously described, *q* can be used to model imperfect sampling for both RT and sequencing.

**Figure 2:**
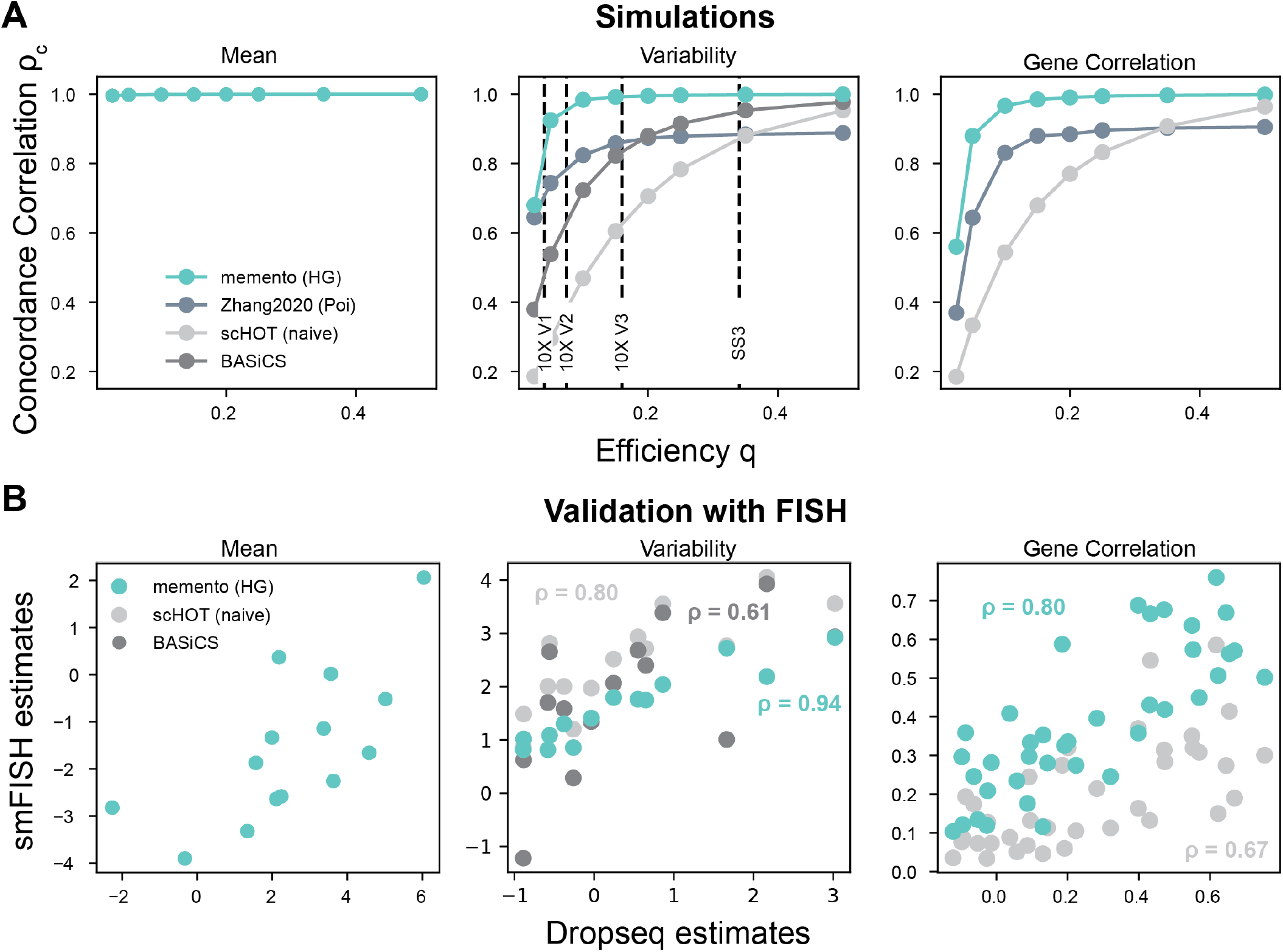
Performance of method of moments estimators. **(A)** Concordance of estimates of mean (left), variability (middle) and gene correlation (right) from scHOT (naive), Poisson, BASiCS and memento with simulated ground truth values (y-axis) for a range of overall transcript capture efficiencies (*q*) (x-axis). Concordance is measured with Lin’s concordance correlation coefficient. **(B)** Scatterplot of estimates of mean (left), variability (middle) and gene correlation (right) of DropSeq data using scHOT (naive), BASiCs, and memento(x-axis) against smFISH-derived estimates measured in the same population of melanoma cells (y-axis). Correlation is measured by Pearson correlation coefficient.

To further assess the accuracy of `memento`’s parameter estimates, we reanalyzed a dataset containing paired droplet-based scRNA-seq and RNA FISH data [38]. This data was previously analyzed using SAVER [39], an imputation method that borrows information from similar genes and cells that has been shown to outperform other approaches including MAGIC and scImpute (**Fig. 2B**). For 16 genes profiled using both scRNA-seq and FISH, the mean estimates from the naive estimator, BASiCS, and `memento` were essentially the same as expected and observed in simulation. For residual variance and gene correlation, estimates produced by `memento` were more correlated with those obtained by FISH (residual variance: *ρ* = 0.94, gene correlation: *ρ* = 0.80) than the naive estimator (residual variance: *ρ* = 0.80, gene correlation: *ρ* = 0.67) and BASiCS (residual variance: *ρ* = 0.61). Importantly, `memento` produces better estimates of residual variance and gene correlation than SAVER without utilizing additional data required by imputation (**Fig. S2**). This advantage results in computational efficiency of estimation (`memento`: 17 seconds vs SAVER: 30 minutes for 16 gene pairs) but also produces estimates that may be better suited for certain downstream analyses (e.g., genetic mapping) where imputation could induce confounding effects due to borrowing information from other genes and cells. These results demonstrate the accuracy of `memento`’s parameter estimates through simulations and comparisons to gold-standard FISH data.

### Hypothesis testing using highly efficient bootstrapping

The goal for hypothesis testing is to determine if an observed difference in the estimated parameters between groups of cells, for example mean, variability and gene correlation, is statistically significant compared to a null hypothesis. When thousands of genes are tested, as is often the case for scRNA-seq experiments that profile the entire transcriptome, we are primarily concerned about the multiple testing problem of nominating a set of candidate genes that can be practically followed up experimentally with an expected number of validations. It is therefore important that the distribution of test statistics under the null hypothesis to be well calibrated amenable to multiple testing correction. While the method of moments estimates using the hypergeometric model are simple to compute and flexible to the true distribution of gene expression within cells, computing confidence intervals (CIs) around the estimates and establishing statistical significance require bootstrapping the data. Bootstrapping large number of cells using a standard scheme that samples cells with replacement would require extensive computational resources that are both time and memory prohibitive, especially for large datasets.

In `memento`, we implemented an innovative scheme that exploits the sparsity and discreteness of scRNA-seq data to enable fast, low-memory, and highly parallelizable bootstrapping. Key to our scheme is the insight that the number of unique observed transcript counts is much smaller than the number of cells (**Fig. S3**). Thus for each bootstrap iteration, instead of resampling individual cells’ counts from a multinomial distribution containing *N* elements (cells) (Multinomial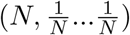 [40]), we only need to resample *K* unique transcript counts for each gene from Multinomial 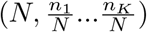, in proportion to the observed frequency of each count (**Fig. 3A**). This amounts to fitting a very small weighted dataset (*K* << *N*) for each resampling iteration. To accommodate multiplexed experiments, we extend our boostrapping strategy using a meta-regression framework that treats each replicate as a separate subgroup of the data to enable hierarchical resampling. In simulation, `memento`’s bootstrapping strategy produces very accurate estimates of the null distribution for mean, residual variance, and gene correlation compared to those obtained with naive resampling (**Fig. S4**). Using bootstrap to quantify the CI in the parameter estimates, `memento` computes well-calibrated empirical p-values for DM, DV and DC that is appropriate for multiple testing correction (**Fig. 3B**).

**Figure 3:**
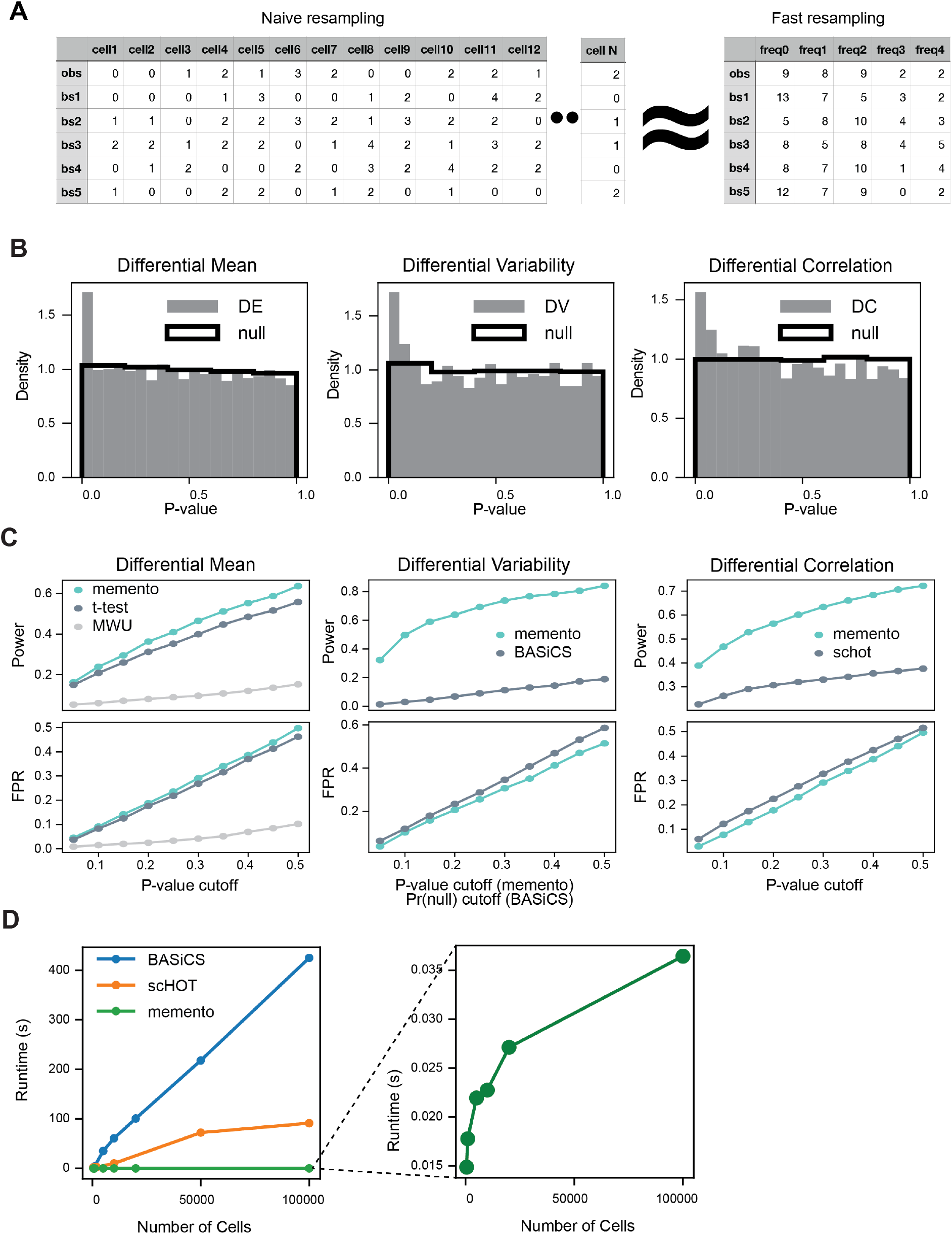
Inference via efficient bootstrapping leveraging data sparsity. **(A)** Schematic for efficient approximate bootstrapping utilizing sparse scRNA-seq data. Rows represent bootstraped samples. The number of unique transcript counts is far smaller than the number of cells. **(B)** P-value histograms from simulated data for differential (DM), variability (DV), and correlation (DC) (left to right). Black line indicates the null distribution where true parameters are held constant across groups. **(C)** Statistical power and FPR (y-axis) against p-value thresholds in simulation for DM, DV, and DC analyses. memento, t-test, and Mann-Whitney U test was used for DM analysis, memento and BASiCS for DV analysis, and memento and scHOT (Spearman r) for DC analysis. **(D)** Simulated runtimes for DV/DC analysis using BASiCS, scHOT and memento(y-axis) for a single bootstrap resampling against the number of cells for binary comparisons (x-axis).

To show that `memento` produces well-calibrated p-values while maintaining high statistical power, we simulated a dataset containing two distinct cell populations. To maintain relevance to real data, we utilized parameters extracted from a real dataset of helper T cells before and after stimulation with rIFNB. We generated a dataset where the estimated differences in the mean, variability, and correlation were maintained for 150 genes and removed for the rest of the genes (see **Methods**). We show that for DM, DV, and DC, `memento` produces well calibrated p-values with the expected number of false positives at a given significance cutoff, while achieving the highest power for detecting true differences (**Fig. 3C**). In particular, for DV and DC tasks, `memento` dramatically outperforms competing methods in power while maintaining a lower false positive rate at each significance threshold.

When compared to existing methods for DM, DV and DC, `memento` is able to perform hypothesis testing orders of magnitude faster, scaling to millions of cells (**Fig. 3D**). In a scenario simulating the throughput similar to emerging scRNA-seq datasets - two groups each containing 10^6^ cells - performing DM and DV analysis for 1,000 genes using 10,000 bootstrapping iterations per gene between groups took 13 minutes using a single CPU. A multicore implementation of `memento` allowed for parallelization of multiple genes, reducing the runtime to 2-3 minutes with 6 CPUs. Particularly for DV and DC, `memento` achieves up to 1000x gain in computational speed given the same resources compared to existing methods. These results demonstrate that `memento`’s bootstrapping strategy produces accurate confidence interval estimates for the effect size at high computational efficiency. These advances results in well-calibrated test statistics and enables hypothesis testing of scRNA-seq data scalable groups of millions of cells (see **Methods** for detailed description of the resampling strategy and hypothesis testing).

### Differential variability and gene correlation in response to exogenous interferon

Interferons are potent immune modulatory cytokines that promote antiviral immunity but associate with the pathogenesis of inflammatory and autoimmune diseases [41]. While interferons are known to act through autocrine and paracrine signaling to induce gene expression, the heterogeneity of the transcriptomic response in stimulated cells has not been extensively characterized. We applied `memento` to investigate how interferon stimulation affects the distribution of gene expression in human tracheal epithelial cells (HTECs). We used multiplexed single-cell RNA-sequencing to profile 69,958 HTECs from two healthy donors across five conditions including unstimulated control and stimulation with type-1 (IFN-*α*, IFN-*β*), type-2 (IFN-*γ*) or type-3 (IFN-*λ*) interferon. For each stimulus, cells were profiled at 3, 6, 9, 24, and 48 hours post stimulation. Following dimensionality reduction, nearest neighbor identification, and Leiden clustering, 7 cell types were identified and visualized using uniform manifold approximation and projection (UMAP) including neuroendocrine cells, iononcytes, tuft cells, basal cells, basal/club cells, globlet cells, and ciliated cells (**Fig. 4A**). Because ciliated cells are known to be the primary target of viral infections including SARS-CoV2 and produce a robust interferon response [42, 43, 44], they were the focus of further analysis.

**Figure 4:**
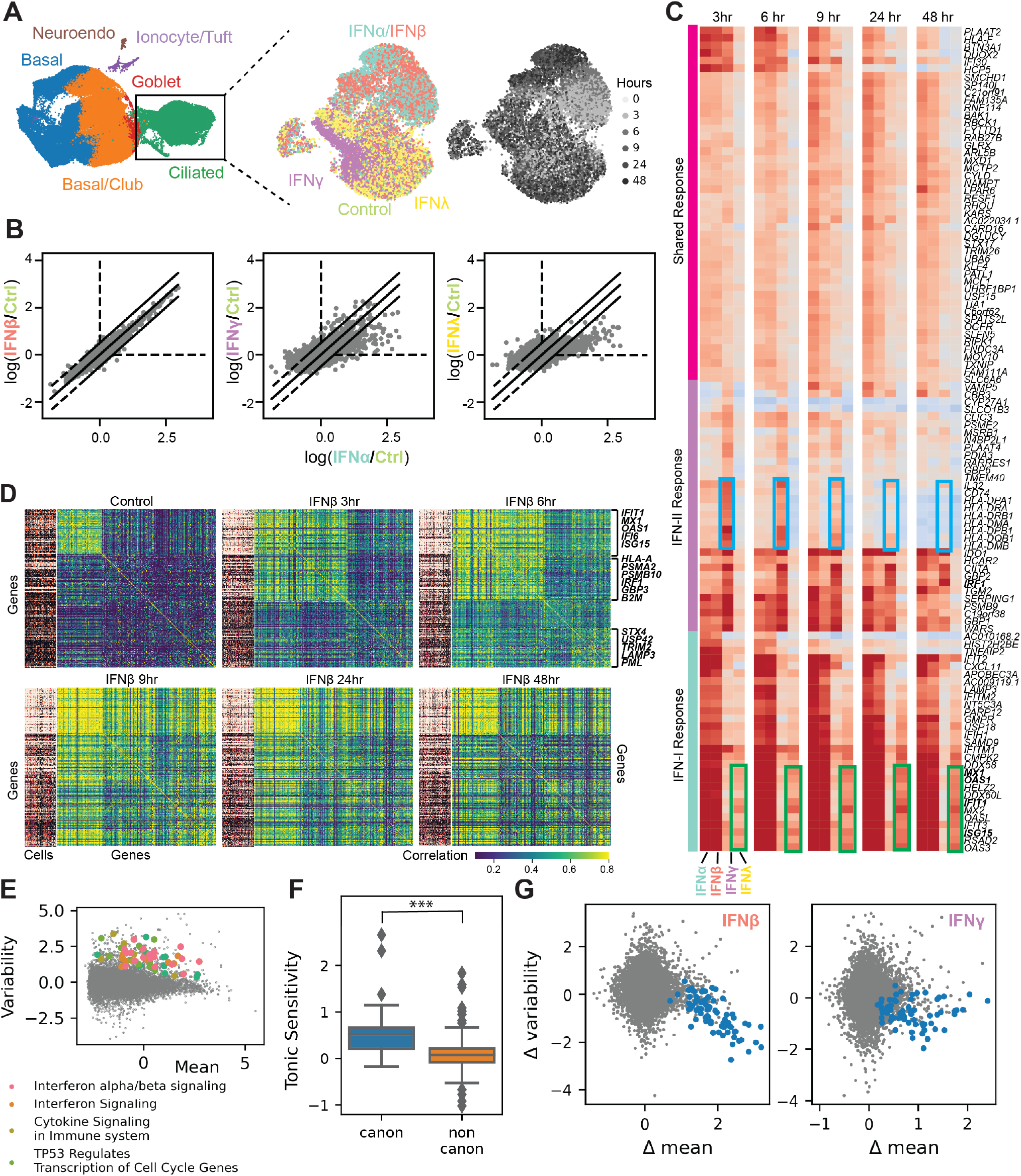
Mapping transcriptional response of human bronchial epithelial cells to extracellular interferon using `memento` **(A)** UMAP of the HBEC dataset with identified cell types (left). UMAP of the ciliated cells (our focus) with stimulation and time labels (right). **(B)** Log fold-change (LFC) of mean expression in response to IFN-*α* (x-axis) against LFC in response to IFN-*β* (left), IFN-*γ* (middle), and IFN-*α* (right) after 6 hours. **(C)** Hierarchically clustered heatmaps of LFC in response to the four types of interferons (columns within each heatmap) across 5 timepoints compared to control. Light blue and green boxes highlight IFN type specific responses. **(D)** Gene-cell heatmap (left of each timepoint) and gene-gene correlation heatmap (right) for selected DMGs in response to IFNb. All genes shown are DMGs detected in response to IFN-*β* at at least one timepoint. **(E)** Baseline expression variability (y-axis) versus mean (x-axis) in ciliated cells. **(F)** Tonic sensitivity (y-axis) for canonical and non-canonical ISGs (x-axis). *** indicates *P* < 0.001. **(G)** Change in variability (y-axis) against the change in the mean (x-axis) for IFN-*β* (left) and IFN-*γ* (right).Blue dots represent canonical ISGs.

We identified 5,018 genes exhibiting differential mean expression (DMGs, FDR < 0.01) between unstimulated ciliated cells and those stimulated by any of four interferons at 6 hours. Comparing IFN-*α* to IFN-*β* and IFN-*λ* revealed similar effect sizes for changes in mean abundance (*ρ* = 0.96), while comparisons to IFN-*γ* identified both type-1 and type-2 interferon specific genes that changed in mean abundance (*ρ* = 0.70, **Fig. 4B**). We define genes that are upregulated in response to interferon as interferon-stimulated genes (ISGs). Hierarchical clustering of the 6-hr ISGs revealed dynamic transcriptomic response shared across the interferons including early induction of MHC class II genes and a cluster of genes consisting of *PLAAT2, BTN3A1*, and *DUOX2* (**Fig. 4C**). We also identified patterns specific to each interferon, exemplified by a subset of canonical ISGs (*IFI2, IFITM2*, and *ISG15*) whose mean expression show a continuous increase in response to IFN-*λ* but remains high for type-1 interferons throughout the time course (**Fig. 4C**).

Analysis of differential mean expression revealed the induction of canonical and non-canonical ISGs (e.g., components of the proteosome) but did not reveal whether these genes are subjected to the same transcriptional regulatory control. To map the interferon gene correlation network and its sub-components, we used `memento` to estimate and compare correlations between pairs of ISGs across stimulation and time (**Fig. 4D**). Agglomerative clustering of the resulting gene correlation matrix revealed distinct subsets of ISGs in response to IFN-*β* that form clusters in unstimulated cells, stimulated cells, or both which could not be distinguished by differential mean analysis. For example, canonical ISGs including *MX1, OAS1*, and *IFI6*, were highly correlated even in the absence of exogenous interferons (**Fig. 4D**). Upon IFN-*β* stimulation, the correlation network consisting of canonical ISGs was expanded to include non-canonical ISGs such as the MHC Class I molecules and other genes associated with antigen presentation that were not correlated in unstimulated cells (**Fig. 4D**). Indeed, more differentially correlated pairs of genes (DCGs, FDR < 0.1) were found among non-canonical ISGs (860 DCGs, 34% of total pairs) than canonical ISGs (421 DCGs, 16% of total pairs). Distinct additional clusters of correlated genes arose as well, such as a cluster at 6hrs that include *STX6, PML*, and *LAMP3*.

We hypothesized that canonical ISGs are correlated in unstimulated cells because of the sensing of tonic interferon and coordinated induction of ISGs by a small number of cells. Tonic interferon signaling has been described to induce a natural gradient of ISG expression across cells [45, 46], which has been further shown to be important for viral defense [46], immune cell homeostasis, and autoimmunity [45]. In our data, canonical ISGs are more variable than non-canonical ISGs in unstimulated cells (**Fig. 4E**), comparable to previously reported differences between cytokines and non-cytokines (**Fig. S5**) [47]. Among the 761 differentially variable genes (DVGs, FDR < 0.1) identified using `memento` between unstimulated ciliated cells and those stimulated by any of the four interferons at 6 hours, 394 were highly variable in unstimulated cells (FDR < 0.005) and were enriched for ISGs (GSEA Interferon alpha/beta signaling Adjusted *P* = 3.35 | 10^−12^) including *IFIT1, IFIT3*, and *MX1*. We next compared the tonic sensitivity of canonical and non-canonical ISGs, estimated as the fold-change (FC) in the expression of each gene between macrophages from *IFNAR* knockout and wild-type mice in the absence of exogenous interferon [48]. This analysis revealed that the canonical ISGs are significantly more sensitive to tonic interferon than non-canonical ISGs (*P* < 2.73 × 10^−10^), **Fig. 4F**). When cells are stimulated with IFN-*β* and to a lesser extent with IFN-*γ*, the variability of many (78% and 39%, respectively) canonical ISGs decreases significantly (**Fig. 4G**, FDR < 0.1), suggesting that exogenous stimulation may homogenize the cellular environment and remove the effects of heterogeneous response to tonic interferon.

These results demonstrate the applicability of `memento` for the comparison of gene expression distributions to reveal novel modes of gene regulation imparted by extracellular interferon. In HTECs, we identified: 1) a core network of canonical ISGs that exhibit highly variable and correlated gene expression in unstimulated cells due to tonic interferon signaling, 2) the synchronization of canonical ISGs in response to exogenous interferon resulting in reduced variability, and 3) a network of non-canonical ISGs that are regulated only in response to exogenous and not tonic interferon.

### Differential expression analysis of perturbed CD4^+^ T cells maps gene regulatory networks in T cell activation

The integration of precise genome perturbations delivered by CRISPR-Cas9 with scRNA-seq profiling of perturbed cells has created new opportunities for forward genetics screens across a variety of in vitro systems. We next applied `memento` to analyze 173,000 CD4^+^ T cells perturbed by CRISPR-Cas9 to map transcriptional regulatory networks that control the activation and polarization of human CD4^+^ T cells. CD4^+^ cells were perturbed using pooled sgRNA lentiviral infection with Cas9 protein electroporation (SLICE) [13] followed by multiplexed single-cell RNA-sequencing (mux-seq). A total of 280 sgRNAs were used in the transduction targeting 140 transcriptional regulators (TRs) that were either highly expressed (top quartile from bulk RNA-seq) or have binding sites that were differentially accessible (from bulk ATAC-seq) in activated CD4^+^ T cells [49] (**Fig. 5A**). Following Cas9 electroporation and multiple rounds of selection and proliferation, activated CD4^+^ T cells from 9 donors were profiled using mux-seq.

**Figure 5:**
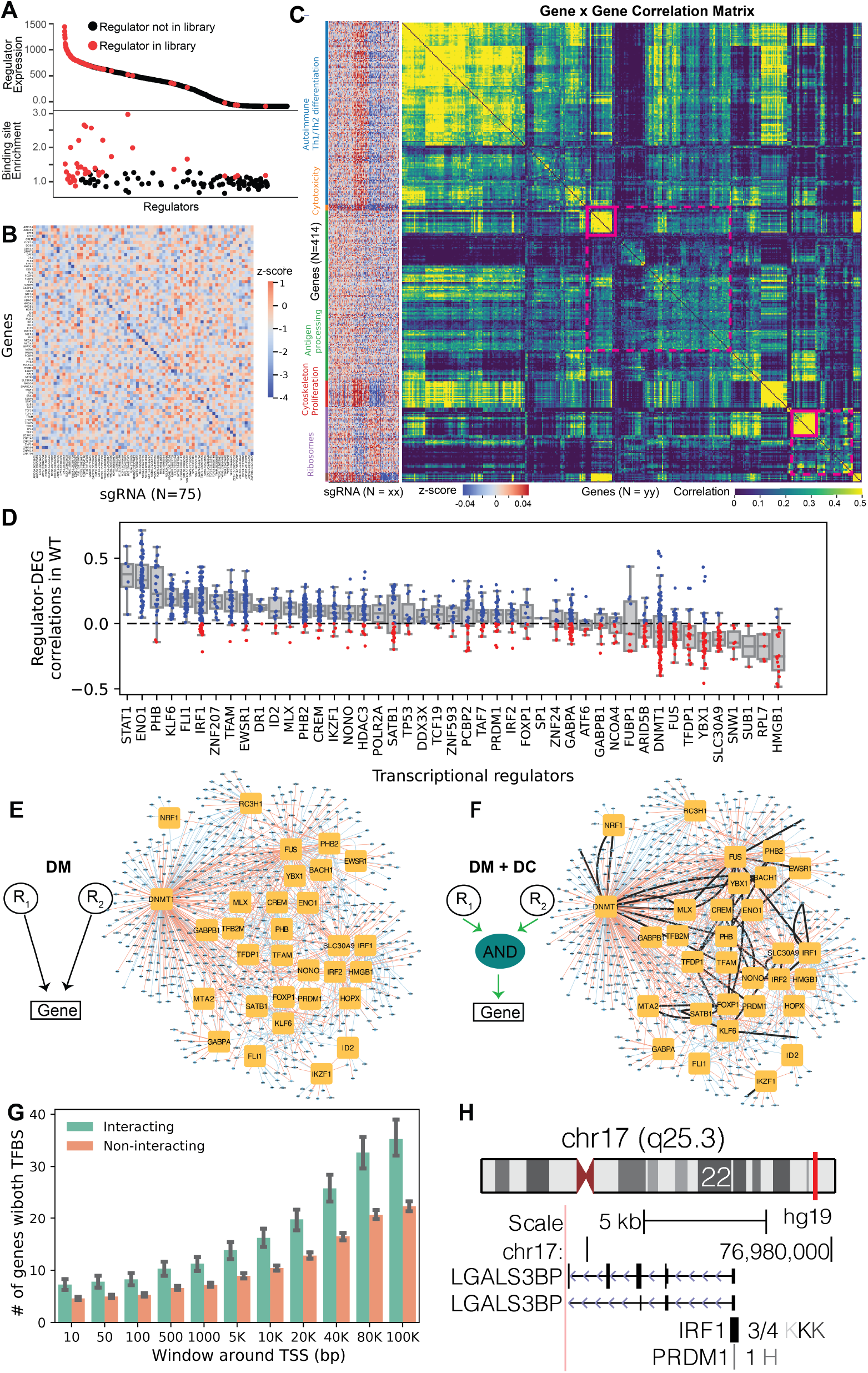
Reconstructing gene regulatory networks of T cell activation using Perturb-seq and memento. **(A)** Selection criteria for perturbed regulators in this study, based on expression (top) and binding site enrichment (bottom). **(B)** Heatmap of gene (row) expression for target genes of sgRNA (columns). **(C)** Left: Heatmap of genes with differential mean expression (row) for sgRNA (columns). Right: Gene-gene correlation matrix for the same DMGs estimated from WT cells. **(D)** Correlation between each regulator and its downstream genes in WT cells. **(E)** Bipartite gene regulatory network that do not account for interaction between regulators constructed from DM analysis of Perturb-Seq data. **(F)** Gene regulatory network including genetic interactions between regulators constructed utilizing both DM and DC analysis. **(G)** Number of genes with binding sites for both pair of interacting or non-interacting regulators across varying windows of the TSS. **(H)** Chromosomal location of *LGALS3BP* and binding sites for IRF1 and PRDM1, predicted to interact using DM and DC analysis.

To assess the cutting efficiency of each sgRNA, we sequenced the sgRNA pool and DNA of edited cells from each donor by targeted amplification of 268/280 loci. Across 268 sgRNAs, the average cutting efficiency, defined as the coverage of edited cells at the target locus divided by the coverage of the corresponding sgRNA in the pool, was 21% (standard deviation 15%, **Fig. S6**). We defined 14 sgRNAs with cutting efficiencies less than 2.0% (standard deviation 1.7%; z-score, *P* < 0.05) as uncut negative controls (WT). We demonstrate the robustness and performance of our screen by the following two quality control assessments. First, we showed using `memento` that target genes were significantly down regulated in cells transduced with the concomitant sgRNA (**Fig. 5B**). Second, there was a higher correlation of average gene expression between WT cells (*ρ* = 0.50) or cells transduced with sgRNAs targeting the same gene (*ρ* = 0.44) than cells transduced with sgRNAs for two random genes (*ρ* = 0; KS-test *P <* 2.2 × 10^−16^ for both; **Fig. S7**).

Using `memento`, we identified 7641 genes (FDR < 0.05) exhibiting differential mean expression (DMGs) between WT cells and cells perturbed by at least one sgRNA. Hierarchical clustering revealed groups of sgRNAs that had similar effects on the transcriptome as well as groups of genes that responded similarly to those perturbations (**Fig. 5C**). Specifically, we identified 5 clusters of DMGs associated with ribosomes (FDR < 5.35 × 10^−24^), cytotoxicity (FDR < 0.014), antigen presentation (FDR < 0.0011), and proliferation (FDR < 0.001). Additionally, the pairwise correlation matrix of DMGs computed using `memento` revealed additional sub-clusters within each of the 5 clusters of DMGs in both WT and perturbed cells (**Fig. 5C**). For example, while the mean expression of antigen processing genes are affected by a common set of transcriptional regulators, a subset of MHC class II genes (e.g., *HLA-DPA1, HLA-DRA, HLA-DRB1, HLA-DPB1*) are highly correlated suggesting that their regulation may be controlled by additional *trans* regulators.

Next, we hypothesized that by leveraging `memento`’s ability to detect changes in gene correlations, we can identify genetic interactions between transcriptional regulators without the need for inducing combinatorial perturbations. We focused the genetic interaction analysis on pairs of DMGs and their transcriptional activators, defined as regulators where a knockout results in the decreased expression of DMGs (TR-DMG, **Methods**). These TR-DMGs tend to be positively correlated with each other in WT cells (Binomial test, *P* < 0.00668, **Fig. 5D**). In the absence of an interaction, two transcriptional regulators (R1 and R2) independently regulate the target gene (G), and knocking out one regulator would not affect the function of the other (**Fig. 5E**). In the presence of an interaction, knocking out one regulator (e.g., R1) would affect R2’s ability to regulate G, which we can detect as a change in the gene correlation between R2 and G when R1 is perturbed (**Fig. 5F**). Following this approach, we identified 564 genetic interactions between 432 unique pairs of regulators (FDR < 0.1, **Fig. 5F**). Validating these interactions, analyses integrating ChIP-seq data from ENCODE[50] show that interacting TR pairs have more target genes with co-localized binding sites near transcription start site (TSS) than non-interaction pairs (**Fig. 5G**). As an example, we identified that *IRF1* regulates *LGALS3PB* (using differential mean expression testing) and the two genes are highly correlated in WT cells (*ρ*_*W T*_ = 0.28). Knocking out of *PRDM1* significantly reduced the correlation between *IRF1* and *LGALS3PB* (Δ*ρ* = -0.38) suggesting that *PRDM1* and *IRF1* may interact to regulate the expression of *LGALS3PB*. Consistent with these observations, *LGALS3BP* has binding sites for both *IRF1* and *PRDMB1* immediately surrounding its TSS (**Fig. 5H**).

These results demonstrate that when paired with a forward-genetic screen such as Perturb-seq, correlation analysis using `memento` can identify sets of genes that share common regulators but may be in different pathways and reconstruct genetic interactions of *trans* regulators that control T cell activation.

### Genetic analysis of population-scale single-cell RNA-sequencing

The growing availability of population-scale scRNA-seq datasets has enabled mapping of genetic variants associated with changes in the expression distribution of proximal genes (*cis*) in specific cell types. Most studies currently deploy pseudobulk methods such as Matrix eQTL to identify *cis* expression quantitative trait loci (*cis*-eQTLs that affect mean expression. While linear mixed models have recently been applied to map *cis*-eQTLs from scRNA-seq data, they are computationally inefficient, limited to the comparisons of means, and sensitive to the underlying parametric model [51]. We hypothesize that compared to pseudobulk methods, `memento`’s more accurate parameter estimates and ability to account for inter-individual variation should increase the power to detect *cis*-eQTLs and discover novel variability and correlation QTLs (vQTL and cQTL, respectively). Furthermore, the highly efficient hierarchical bootstrapping strategy should enable applications to the largest population-scale scRNA-seq datasets that may be computationally prohibitive for complex parametric linear mixed models. To demonstrate, we applied `memento` to reanalyze a previously published scRNA-seq dataset containing 1.2M PBMCs from 160 SLE patients and 90 healthy donors.

The data was analyzed separately for each of the following reported cell types: CD4 T cells (T4), CD8 T cells (T8), natural killer cells (NK), classical monocytes (cM), and non-classical monocytes (ncM) [17]. Individuals of East Asian and European ancestries were separately analyzed to enable replication analysis of `memento` by comparing the results between populations. For each cell type and ancestry, `memento` mapped *cis* genetic variants (e.g., within 100kB from the TSS) associated with expression mean, expression variability, and gene correlation with well-calibrated p-values (**Fig. 6A**). We compared the power and false positive rate (FPR) of `memento` and Matrix eQTL for detecting *cis*-eQTLs reported by the OneK1K study consisting of 1000 non-overlapping individuals [18]. In both East Asians and Europeans, `memento` (AUC=0.85) had more power to detect *cis*-eQTLs compared to Matrix eQTL (AUC=0.81) given the same FPR (**Fig. 6A**,**B**). Overall, `memento` outperformed Matrix eQTL in both populations, replicating 1,606 vs 855 *cis*-eQTLs across cell types in East Asians and 1,778 vs 958 in Europeans. Further, across a range of individuals representative of existing cohort sizes for multiplexed scRNA-seq experiments, `memento` achieved an average gain in power of 15% for 80 individuals that increased to 32% for 50 individuals with an average of 440 cells per individual (**Fig. 6B**).

**Figure 6:**
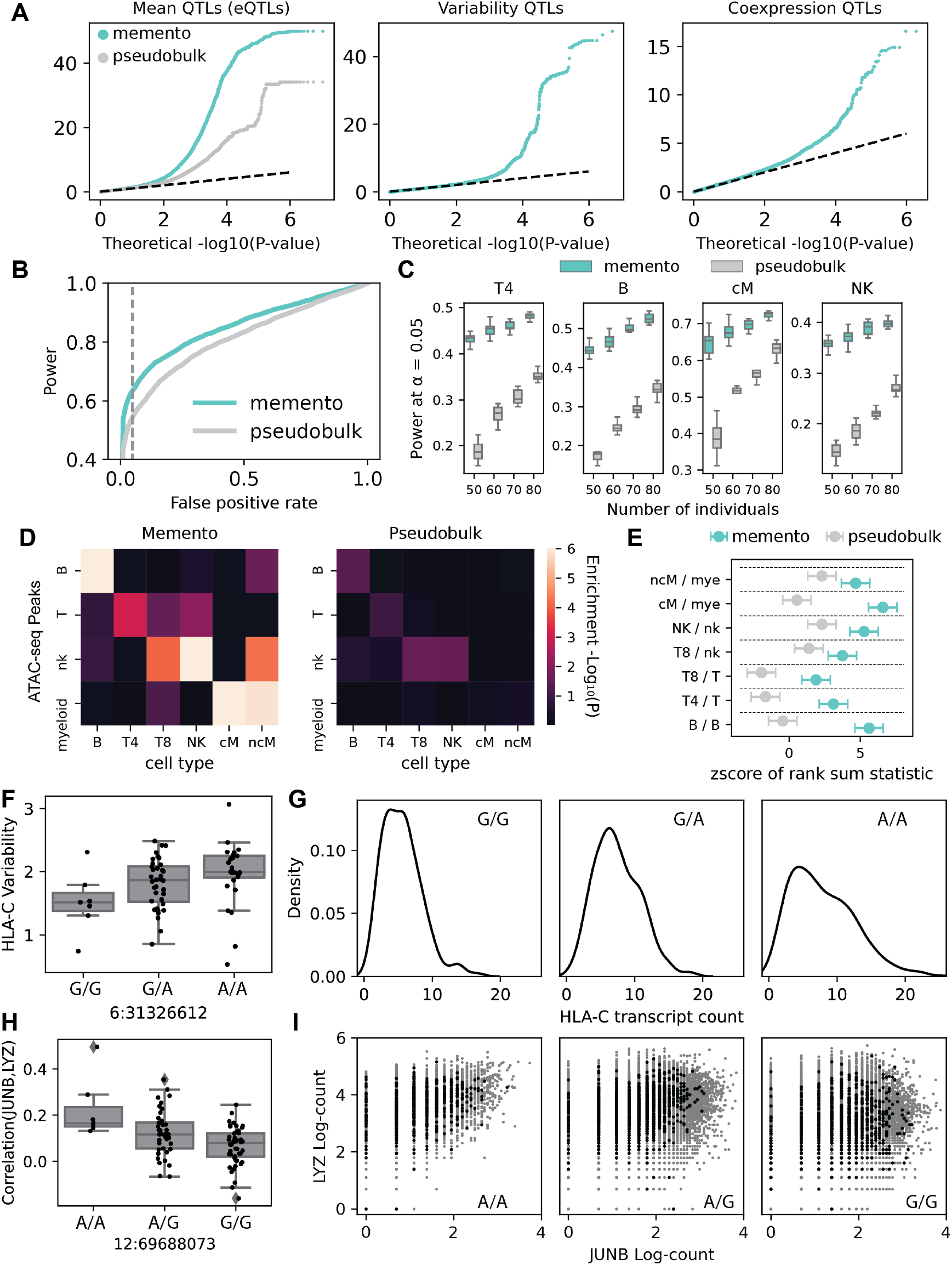
Mapping of mean QTL (eQTL), variability QTL (vQTL), and correlation QTL (cQTL) using memento. **(A)** Quantile-quantile (QQ) plots for expected p-values (y-axis) computed by memento versus theoretical p-values (x-axis) for mean, variability, and correlation QTLs. For mean QTLs, QQ-plot of p-values from pseudobulk approach (Matrix eQTL) is overlayed. **(B)** Receiver operating characteristic (ROC) curve for recovery of mean QTLs identified from a much larger cohort (OneK1K) for memento and pseudobulk method. **(C)** Power of eQTL recovery (y-axis) of memento and pseudobulk method across different numbers of individuals. Analyses was performed on CD4 (T4), B cells (B), classical monocytes (cMs) and natural killer cells (NKs). **(D)** Enrichment of cell-type specific eQTLs in cell-type specific ATAC-peaks. Each entry represents the enrichment for eQTLs detected in one cell type (column) in ATAC-peaks detected in another cell type (row). Intensity is log10(p-value). **(E)** Enrichment of eQTLs detected in each cell type for cell-type-specific ATAC-peaks detected in the same cell type. **(F)** An example of a variability QTL. Expression variability (y-axis) for each individual of varying genotypes at6:31326612. **(G)** Histogram showing distribution of *HLA-C* expression for a representative individual of each genotype. **(H)**An example of a correlation QTL. *JUNB* -*LYZ* gene correlation (y-axis) for individuals of varying genotypes at 12:69688073.**(I)** Scatterplot of expression of *LYZ* (y-axis) against the expression of *JUNB* (x-axis) across single cells from all donors (grey) and a representative individual (black).

We next investigated whether the increased number of *cis*-eQTLs detected by `memento` also improves the enrichment for regions of open chromatin and disease associations. In the East Asian population, *cis*-eQTLs identified by `memento` in specific cell types were more enriched for cell type specific regions of open chromatin annotated by an independent study that performed ATAC-seq on bulk sorted immune cells (p-values for matched cell-types, B 9.0×10^−9^ vs 0.04; T4 9.3×10^−4^ vs 0.11; T8 0.03 vs 0.58; NK 6.67×10^−8^ vs 0.03; cM 2.1×10^−11^ vs 0.67; ncM 1.0×10^−6^ vs 0.46, **Fig. 6D**,**E**). Similar gains in enrichment was observed in the European population (**Fig. S8**). Analysis using LD score regression found that *cis*-eQTL identified by `memento` were also more enriched for GWAS associations to immune mediated diseases suggesting improved performance for fine mapping (**Fig. S9**).

Beyond mapping cis-eQTLs, `memento` enables mapping of genetic variants associated with expression variability and gene correlation that can suggest additional mechanisms by which genetic variants affect gene expression. Using `memento`, we identified 10607 expression variability QTLs (vQTLs) for 733 genes across all cell types. For example, the expression variability of *HLA-C* differs between different genotypes of 6:31326612 (**Fig. 6F**). The A allele increases the expression variability of *HLA-C* without a significant effect on the mean (**Fig. 6G**). For mapping correlation QTLs (cQTLs), we focused on testing for the correlation between genes with at least one significant *cis*-eQTL and known transcription factors. This choice specifically tests the hypothesis that the genetic variants may further modify the effect of transcription factors on gene expression. We mapped 3726 cQTLs for 238 pairs of genes across all cell types. For example, the SNP at 12:69688073 not only affected the mean expression of *LYZ*, but also the the correlation between *JUNB* and *LYZ*. Interestingly, there exists a JUNB binding site within 1kbp of the SNP suggesting that JUNB may act as a *trans* regulator for *LYZ*, and that its strength of regulation is affected by the genotype at this site.

These results demonstrate `memento` as a scalable method for the genetic analyses of population scale scRNA-seq data delivering higher statistical power for the identification of *cis*-eQTLs and new capability for mapping variability QTLs and correlation QTLs. These advances improve the fine mapping of disease associations and reveal novel modes by which genetic variants can affect gene expression.

## Discussion

Fueled by the development of scalable workflows, there is an emergence of scRNA-seq datasets where the quantitative comparison of gene expression distributions between groups of cells is a critical task. These include efforts to characterize the differences in single-cell expression profiles between experimental conditions [12], between cells harboring different genetic perturbations induced by genome editing [14, 52], and between individuals inheriting different alleles [16, 17, 18]. Initial observations from these studies that experimental and genetic perturbations often induce subtle shifts in gene expression rather than distinct cell states have necessitated the need for methods that compare gene expression distributions. However, scalable computational methods that facilitate hypothesis testing over many covariates (e.g. hundreds of *in vitro* perturbations or millions of genetic polymorphisms) are still scarce. Furthermore, even fewer methods currently test for differences in the variability of gene expression and correlation between pairs of genes, parameters that are uniquely captured from single cell RNA-sequencing. Here, we introduced `memento`, an end-to-end method for the quantitative analysis of scRNA-seq data theoretically scalable to millions of cells.

`memento` is developed with two key innovations: method of moments estimators that model scRNA-seq as a hypergeometric sampling process and an efficient bootstrapping strategy to construct accurate confidence intervals around parameter estimates leveraging the sparsity of scRNA-seq data. Method of moments estimators provide two advantages over other approaches. First, our approach explicitly disentangles the biological and technical sources of noise to accurately characterize biological variation. This feature of `memento` addresses recent calls for hierarchical parametric modeling of the measurement noise of scRNA-seq while only considering biological variation for estimation and inference [22]. Second, given a hierarchical model of scRNA-seq, computing the overall likelihood with a discrete component is computationally prohibitive due to the need to marginalize over each possible level of biological gene expression. Method of moment estimators obviate the need to repeatedly compute the overall likelihood and directly computes the parameters of interest instantly. The multinomial approximation of hypergeometric sampling has been used to theoretically derive the baseline noise in scRNA-seq [33] and to design dimensionality reduction techniques for count data [53]. The Poisson approximation of the binomial (which in turn approximates the hypergeometric), has been used to derive empirical Bayes estimators to inform the optimal design of scRNA-seq experiments [36]. While we derive our estimators focusing on scRNA-seq workflows where the cell-to-cell differences in transcript sampling frequencies *q* is small, hypergeometric formulation is flexible to modeling workflows where *q* may differ significantly between compartments (e.g. sci-rna-seq[30]), provided that *N*_*c*_ and *q* can be estimated separately. Because of the modular and flexible nature of `memento`, we further anticipate that our modeling framework could be extended to alternative scRNA-seq workflows that use hybridization instead of reverse transcription [54] and spatial transcriptomics data [55, 56]. Analyses of emerging multimodal workflows (e.g., ATAC-seq and CITE-seq) should also be possible by modifying the method-of-moments estimators to correctly capture sources of technical variation unique to each assay.

The general challenge in implementing method of moments estimators for hierarchical models is the need to establish confidence intervals using resampling because incorporating the sampling process into deriving analytical confidence intervals and p-values are likely to have highly complex forms without further assumptions. While resampling can be computationally prohibitive especially when the number of cells is large, our use of the approximate bootstrap that resamples the number of unique counts rather than number of single cells enables us to adopt a method of moments approach. When subsampled at various cell counts, the number of unique counts increased sub-linearly with the number of cells (**Fig. S3**), and this was true even when considering unique counts for pairs of genes (**Fig. S10**). Through extensive simulations, we demonstrated that `memento` is able to produce accurate confidence intervals for the moment estimates and well-calibrated p-values testing for their differences across groups of cells. Because our hypothesis testing framework utilizes approximate bootstrapping, it should in theory be compatible with existing parametric models to enable better estimates of empirical p-values for a variety of single-cell sequencing analysis methods.

Through three proof of principle applications of `memento`, we demonstrate how differential variability and correlation analysis can identify novel gene regulatory relationships that are not detected using differential mean analysis. In human tracheal epithelial cells, we showed that `memento` identified unexpected correlation of canonical ISGs at baseline suggestive of an extracellular gradient of tonic interferon, and the expansion of the interferon response transcription regulatory network after extracellular stimulation to include non-canonical ISGs. In a dataset of CD4^+^ T cells genetically perturbed by CRISPR-Cas9, `memento` analyses of gene correlations while using genetic perturbations as causal anchors revealed genetic interactions of regulators in controlling the expression of target genes. Finally, when applied to a population-scale scRNA-seq experiment, `memento` improved the statistical power and resolution for mapping *cis*-eQTLs and mapped additional loci that affected gene expression variability and gene correlation. These applications to diverse datasets demonstrate that `memento` is a highly adaptable and scalable method for the quantitative analyses of large scRNA-seq datasets containing many replicates and experimental conditions.

## Methods

### Modeling scRNA-seq as a hypergeometric sampling process

We model the count data obtained from scRNA-seq with a flexible hierarchical model that explicitly considers the generative process of the expressed transcript counts and sampling of mRNA molecules with massively-parallel scRNA-seq methods. As presented in the main text, our full model of the scRNA-seq sampling process can be summarized as follows:

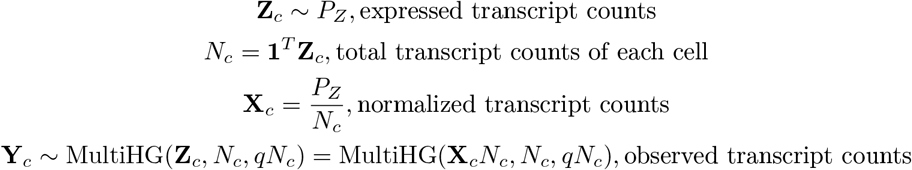

*q* is the random variable representing the proportion of expressed transcript counts that is eventually counted as UMIs in the observed scRNA-seq experiment. In our discussion of sources of noise above as applied to most scRNA-seq workflows, it accounts for both the RT sampling efficiency as well as the sampling of transcripts from sequencing. In the extreme, if a library is sequenced to saturation, then *q* reduces to the RT sampling efficiency; however, in most experiments, libraries are not sequenced to saturation but up to a known percentage of unique molecules. Through extensive simulations, we demonstrate that this compound noise process can be well approximated with a single multivariate hypergeometric process by using a value for 𝔼[*q*] that is a product of the RT sampling efficiency (available for specific experimental technologies) and the sequencing sampling efficiency (available from the preprocessing pipelines such as CellRanger) (**Fig. S1**) [57].

We then model the mRNA capture process with a multivariate hypergeometric distribution. The probability mass function (PMF) of the multivariate hypergeometric distribution given (*K*_1_, *K*_2_, *K*_3_, …*K*_*c*_) components (i.e. genes), total count 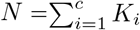, and number of samples *n* ∈ 0, 1, …, *N* is given by:

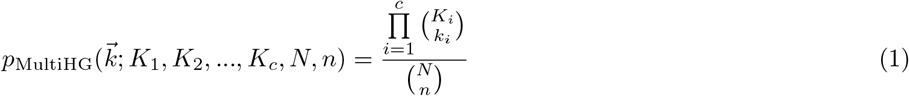

In previous works [36], the full hypergeometric treatment was simplified by a series of approximations, starting from the hypergeometric model to the Poisson model:

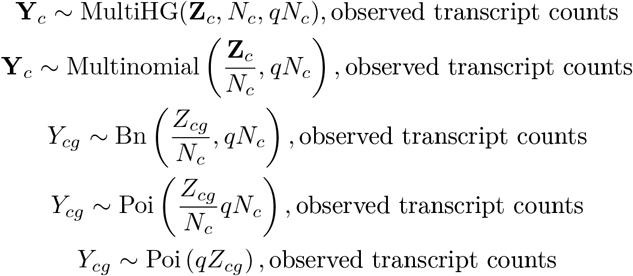

*Y*_*cg*_ is a single element in the vector **Y**_*c*_, as the Poisson model considers the sampling of each gene to be independent. As we discuss in the following sections, the full hypergeometric treatment and the Poisson simplification result in very similar estimators when *q* is very small (close to 0), but become more different as the value of *q* increases, as scRNA-seq experimental workflow improves.

### Method of moments estimation of expressed transcript counts

We will start this section by reviewing the derivation of the Poisson estimators first presented in [36] in the context of determining optimal sequencing depth for scRNA-seq experiments. First, recall the previously presented Poisson sampling model for scRNA-seq where *N*_*c*_ represents the total expressed transcripts for each cell, *q* is the overall sampling efficiency, and *X*_*cg*_ is the true relative mRNA expression *Y*_*cg*_ ∼ Poi (*qN*_*c*_*X*_*cg*_).

For a Poisson variable *A ∼* Poi(*λ*), the moments of *A* are 𝔼[*A*] = *λ* and 𝔼 [*A*^2^] = *λ*^2^ + *λ*. Similarly, for our model, we can write down the equations for the moments of *Y*_*cg*_ given the other variables, *q, N*_*c*_, and *X*_*c*_.

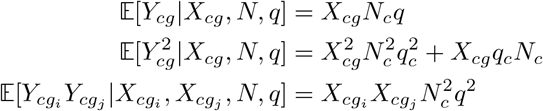

Substituting the first moment equation into the second, we get:

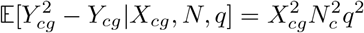

These equations lead to an estimator for 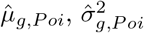, and 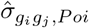, the mean, variance, and covariance of *X*_*cg*_ by averaging the moments over all cells:

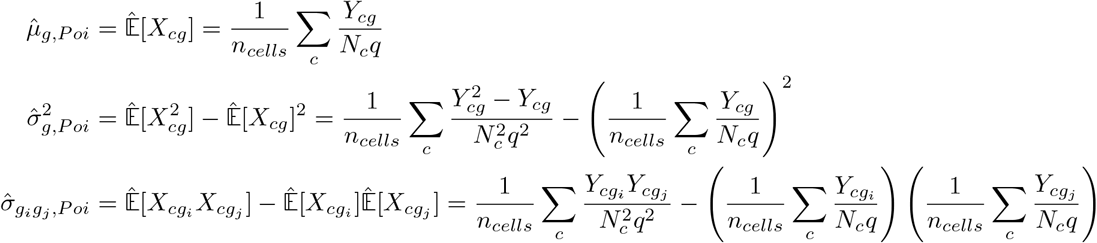

Now, let us consider the full multivariate hypergeometric model, **Y**_*c*_ ∼= MultiHG(**X**_*c*_*N*_*c*_, *N*_*c*_, *qN*_*c*_). For a random vector **A** ∼ *MultiHG*(**K**, *N, n*), the moments of A are:

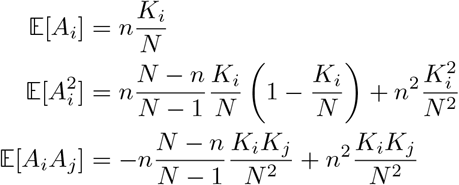

We can again write down the moment equations, this time for the multivariate hypergeometric model.

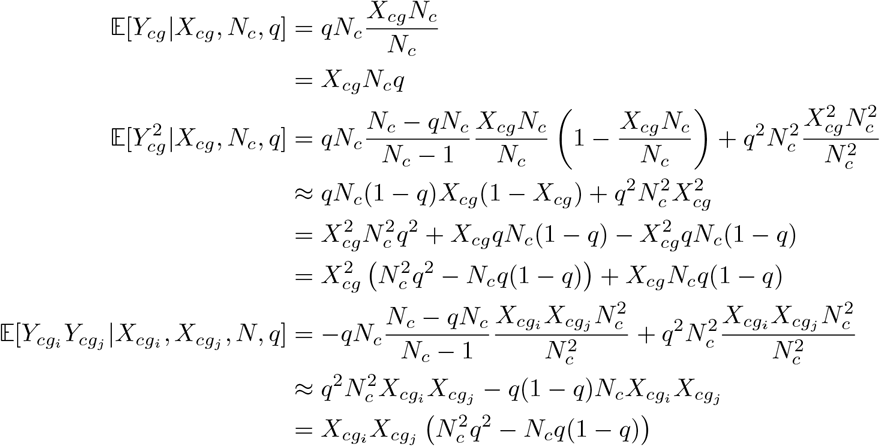

Substituting the first moment equation into the second, we get:

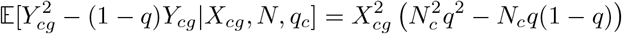

The approximation used in the derivation for the second and first pairwise moment assumes that *N*_*c*_ *>>* 1. For most mammalian cells with expressed transcript counts on the order of 10^5^, these approximation should hold. Similar to estimators based on the Poisson model, we can derive estimators based on these moment equations from the full multivariate hypergeometric model:

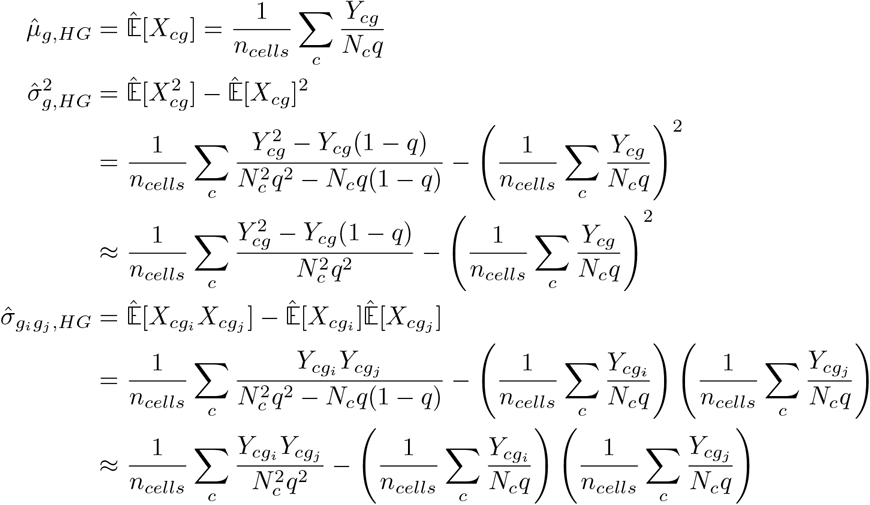

Last, we write the naive estimators for mean, variance and covariance for completeness.

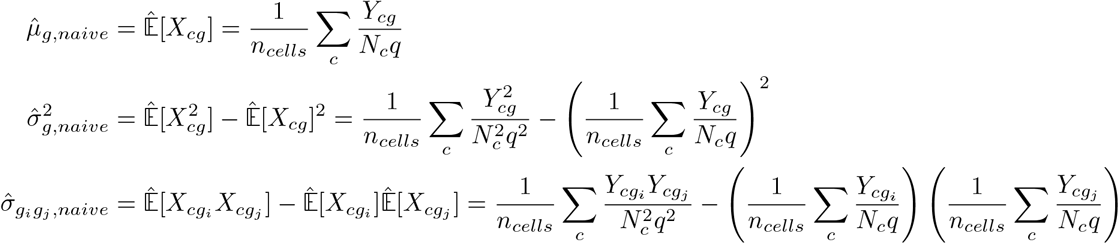

The estimators for the mean and covariance is very similar between the naive, Poisson and HG estimators. However, the estimator for the variance, which contributes to the measurement of residual variance and correlation, is the key difference between the three sets of estimators. Importantly, it is straightforward to see that the HG estimator for the variance includes the naive and Poisson estimators:

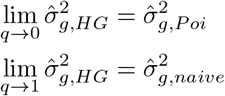

These results imply that when *q*, the overall sampling efficiency, is small, the HG estimators behave very similar to the Poisson estimators. When *q* approaches 1, a hypothetical scenario where the scRNA-seq workflow is perfect and we capture all expressed transcripts, the HG estimators converge to the naive estimator, as there is no noise process. As scRNA-seq workflows improve and *q* becomes larger, HG estimators serve as a generalization of the estimators presented by Zhang et al. to account for different types of experimental workflows with different values of *q*.

We also discuss here the case where *q* is not constant across cells. One of the assumptions used in deriving our estimators above is that *q* is a known constant, and we do not need to estimate it for each and every cell. However, it is plausible that for certain scRNA-seq technologies and when sequencing is not saturated, *q* is actually a distribution around its mean, 𝔼[*q*]. Experimentally, we can mitigate this issue by using spike-in RNA control to actually measure the value of *q* for each and every cell. Because *q* does not appear in the Poisson estimators, it is not possible to explicitly account for the variability in *q* even if its value can be measured for each cell. With the hypergeometric estimators derived here, we can simply substitute the measured values of *q*_*c*_ for each cell in place of *q* above.

### Estimating cell sizes by trimming variable genes

The *N*_*c*_*q*_*c*_ values that appear in the HG estimator equations above refer to the cell size, which serves as a normalization factor for each cell. These constants serve to ensure that even if the proportions of transcripts captured vary across cells, the estimates would not be affected by this technical source of noise. We can decompose *N*_*c*_*q*_*c*_ into two components: a constant *n*_*umi*_ and *γ*_*c*_ so that *N*_*c*_*q*_*c*_ = *n*_*umi*_*γ*_*c*_. The simplest way of estimating *γ*_*c*_ is to first compute 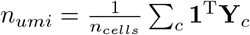, and setting 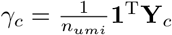, performing a total count normalization.

This is how the Poisson estimators presented in Zhang et al.’s work estimated the cell sizes. In `memento`, we provide an alternate method by first computing residual variances across all cells in a dataset with total count normalization, and trimming off genes that exhibit high variability. This approach assumes that most genes in the dataset should not be differentially expressed, and the least variable genes are appropriate to be used in normalization. This idea of using non-DE genes have been used in other methods, such as [58, 59]. By default, `memento` uses 10% of the least variable genes. After gene set *G∗* is formed by trimming variable genes, we compute *γ*_*c*_ with:

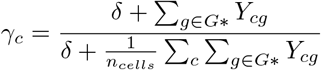

The *δ* value here serves as a regularization factor in estimating cell sizes; when this value is high, it would indicate the dataset does not need a size factor normalization (sampling is truly constant across cells, such as when sequencing to saturation). By default, `memento` uses median(Σ*g*∈*G*∗ *Y*_*cg*_) over cells *c* as the *δ* value.

It is important to note that there are more sophisticated normalization methods that exist in literature [60]. `memento` can readily incorporate these alternative methods of computing cell sizes into its pipeline.

### Computing the residual variance

Mean and variance in scRNA-seq data is generally highly correlated and measuring variability of expression must account for this correlation. BASiCs accounts for this dependence by performing nonlinear regression with many components between the fitted mean and ovedispersion parameters [27]. Instead of fitting a negative binomial distribution then regressing out the mean from the overdispersion parameter, we simply take the estimated true mean and variances and fit a simple polynomial regression. We use a single fitted polynomial (default degree 2) for all genes of a given group of cells, defined by cell type, experimental condition, or batch. We find that even this simple regression is able to largely remove the mean-variance dependence present in scRNA-seq data.

### Efficient bootstrapping by exploiting data sparsity

Typically, generating confidence intervals and computing p-values for hypothesis testing make certain assumptions on both the distribution of the data as well as the estimator itself. For example, to compute p-values for the coefficients of a linear regression model, we typically assume that the data is normally distributed and the sampling distribution of the coefficients are also normal. In the setting of scRNA-seq, our estimators allow for measurement of the average, variability, and gene correlation without making any assumptions about the distribution of expressed transcript counts. However, it is difficult to compute analytical confidence intervals for our estimators without assuming anything about the data itself and the sampling distributions of our estimates.

Bootstrapping is a procedure for estimating the sampling distribution of any arbitrary statistic without making large assumptions on the data generating processs [40]. In `memento`, we propose a strategy to perform bootstrapping in scRNA-seq data in an extremely efficient manner. Specifically, in a dataset for a single gene with *N* cells *x*_1_, *x*_2_, *x*_3_, …*x*_*N*_, we can model the number of appearance of each observation as a multinomial distribution with Multinomial 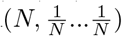. If there are *K* unique counts with *n*_k_ cells each, we can re-write the resampling distribution as Multinomial 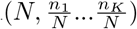.

When considering normalized transcript abundances, we must account for the total number of transcripts in each cell (*N*_*c*_). While this would technically create a different *N*_*c*_ for each cell and make our scheme less useful, a strategy binning *N*_*c*_s across cells into a small number of discrete bins well-approximates the true bootstrap distribution of parameters. Through simulations, we show that as the number of bins increase, we show that the true bootstrap distribution and the approximate bootstrap distributions are nearly identical (**Fig. S4**).

### Hypothesis testing and extension to account for replicates in multiplexed scRNA-seq experiments

Consider a scenario with two groups of cells A and B, and we computed the parameter of interest *t* for each group and computed Δ*t* as their difference. *t* would depend on the type of test we would like to perform; we would compute the mean, residual variance, and correlation to test for differences in the averages, variability, and coexpression respectively. We then perform bootstrapping with *B* iterations to generate a sampling distribution for the test statistic Δ*t*, from Δ*t*_1_ to Δ*t*_*B*_. If we wished to test for the alternative hypothesis of H1: Δ*t* ≠ 0 against the null H0: Δ*t* = 0, we first generate the null distribution by subtracting Δ*t* from Δ*t*_1_, …, Δ*t*_*B*_ to form 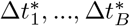, similar to the strategy laid out in [40]. We can then compute the achieved significance level (ASL) for that test as

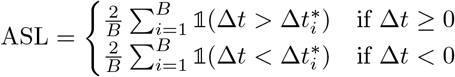

There has been an increasing trend to generate scRNA-seq data with replicates (e.g. different individuals), especially with multiplexed workflows. Consider an experiment with two conditions and *n* replicates. Then, we propose a meta-analysis framework where we first group the cells into 2*n* groups and perform a meta-regression with 2*n* observations:

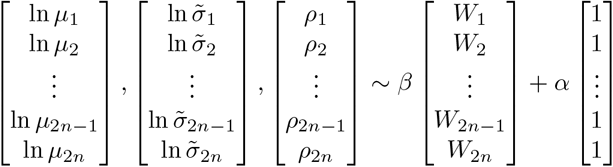

Where 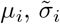, and *ρ*_*i*_ refer to the estimated mean, residual variance, and correlation computed in the *i*^th^ replicate and *W*_*i*_ refers to the condition. Then, we can bootstrap the regression coefficients *B* times to yield the original statistic 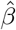 and bootstrap statistics 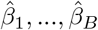. Then, similar to the non-replicated case, we can generate the null distribution 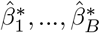, by subtracting 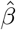 from 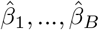,. We can further compute the ASL with:

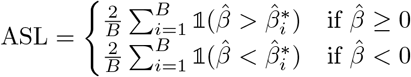

This framework can easily extended to incorporating many covariates, including batch variables and interactions between variables of interest.

As a technical aside, we note that this procedure for computing the ASL assumes that the sampling distribution of the test statistic of interest is translation invariant [40]. Through extensive simulations, we confirm that for the test statistics we consider in `memento`, this procedure yields well-calibrated results under the null hypothesis (**Fig. 3B**). If custom test statistics are used, it is important to check for the calibration of hypothesis test results. `memento` also has the option to compute p-values assuming that the sampling distribution of the effect size is normal with unknown variance that is estimated using the bootstrap, useful for speeding up hypothesis tests. For this work, this approximation was only used for analyzing the effect of natural variation (**Fig. 6**).

### Pre-processing the rIFNB1 PBMC dataset

We used the original clustering and tSNE visualization of the rIFNB1 dataset [15] from the data deposited in the Gene Expression Omnibus under the accession number GSE96583. Further details on the pre-processing of this dataset can be found in the original paper [15].

For all analysis, we selected genes where the mean observed expression 𝔼[*Y*_*cg*_] = 0.07, which was the reliability limit for this experiment. More details on the reliability limit can be found in [36]. This value was computed from the reported UMI capture efficiency of 10X Chromium V1 and well as the sequencing saturation of this experiment, which was around 90% [15].

### Simulating genes with differential mean, variability, and correlation

For simulations presented in **Figure 2** and **3**, we first extracted moments of *X*_*cg*_ and *N*_*c*_ for every gene with E[*Y*_*cg*_] *>* 0.07 from CD4^+^ T cells in the rIFNB1 stimulation data using the HG estimator presented above. We then used these estimated moments to simulate expressed mRNA transcripts by sampling from the negative binomial distributions specified by these moments.

We also induced random correlations between the negative binomial distributions by using the Gaussian copula method. We first sampled a random positive semidefinite matrix using the `scikit-learn` Python package and converted the matrix into a correlation matrix. This correlation matrix was used to generate the Gaussian random variables used in the copula method.

To produce datasets to be compared in 3, we used the procedure above on the control and stimulated cells separately. We then removed the differences for some genes and pairs of genes by replacing the parameters between the groups with their average. These genes produced a ground truth null comparisons by forcing the underlying parameters to be identical, while the other genes served as simulated DEG/DVG/DCGs with a natural distribution of effect sizes.

The simulated ground truth datasets were than subject to hypergeometric sampling at various overall capture efficiencies *q* for For 3, we used *q* = 0.07, corresponding to 10X Chromium V1 chemistry with 50% sequencing saturation.

### HTEC interferon stimulation experiment

Human tracheal epithelial cells were harvested from deceased organ donors according to established protocols (PMID: 1616056). Frozen cell aliquots were reactivated and cultured in epithelial growth media (EGM) [3:1 (v/v) F-12 Nutrient Mixture (Gibco)–Dulbecco’s modified Eagle’s medium (Invitrogen), 5% fetal bovine serum (Gibco), 0.4 ug/mL hydrocortisone (Sigma-Aldrich), 5 ug/mL insulin (Sigma-Aldrich), 8.4 ng/mL cholera toxin (Sigma-Aldrich), 10 ng/mL epidermal growth factor (Invitrogen), 24 ug/mL adenine (Sigma-Aldrich), and 10 uM Y-27632 (Enzo Life Sciences)] on 10 mm dishes coated with rat tail collagen (Sigma-Aldrich). EGM was changed three times a week until dishes were confluent, at which point the cells were passaged with 0.25% trypsin for 30 minutes. For air liquid interface culture, expanded basal cells were plated at 50,000 cells per 6.5 mm transwell insert (Corning 3470) coated with human placental collagen (Sigma-Aldrich) and cultured with Pneumacult ALI (StemCell) for 21-28 days according to the manufacturer’s instructions. Starting on day 27, interferon stimulation (IFN-*β*: 10 ng/ml, IFN-*α*2: 10 ng/ml, IFN-*γ*: 10 ng/ml, IFN-*λ*2: 10 ng/ml) was added at hours 0, 24, 39, 42, and 45 prior to harvesting (For final timepoints 3, 6, 9, 24, and 48 hours). On the day of harvest, basal media was aspirated and both basal and apical chambers were rinsed twice with PBS. Following two washes, trypsin-EDTA (0.25% Fisher cat. 25200072) was added to both the basal and apical chambers (300 ul basal, 100 ul apical) and incubated for 30 minutes at 37°C while pipette mixing every 10 minutes. Trypsinization was quenched with 300 ul of maintenance media and transferred to a 1.5ml eppendorf tube (eppendorf cat. 022431021) and centrifuged at 350xg for 5 minutes at 4°C. Cells were resuspended in 94 ul of cell staining buffer (Biolegend cat. 420201) and blocked with 5 ul of TruStain FcX (Biolegend cat. 422302) for 10 minutes on ice. Blocked cells were stained with 1 ul of Biolegend Totalseq-B hashtags (Biolegend Totalseq-B hashtags 1-11) for 30 minutes on ice. Staining was quenched with 1 ml of cell staining buffer and spun at 300xg for 5 minutes at 4°C prior to two more washes with 1 ml of cell staining buffer. Cells were resuspended in 100 ul of 0.05% BSA in PBS and counted via Countess II (Fisher cat. A27977). Counted cells were pooled equally into two pools and spun at 300xg for 5 minutes at 4°C. Cells were strained through a 100 *µ*M filter (Corning cat. 431752) prior to a final count and each pool was loaded onto two 10x 3’v3 lanes. Libraries were prepared as described in the 10x 3’v3 user guide. Samples were sequenced on three lanes of NovaSeq S4.

### Clustering the HTEC transcriptomes

We performed filtering, normalization, and clustering with the scanpy [31] suite of tools using the default values. Cell types were manually identified based on previously known marker genes for HTECs [44].

Similar to the rIFNB1 dataset, we selected genes where the mean observed expression 𝔼[*Y*_*cg*_] = 0.07, which was the reliability limit for this experiment.

### Clustering the correlation matrices for genes with differential mean expression

DMGs in ciliated cells were identified by using `memento` by comparing each stimulation and timepoint to the unstimulated control. The correlation between the DMGs were computed using `memento` for each timepoint in IFN-*β* stimulation condition. This correlation matrix at timepoint 6hr was then clustered using the AgglomerativeClustering function in sklearn python package. Top 4 clusters in terms of gene number were chosen for plotting.

### Identifying highly variable genes at baseline

We used `memento` in the one-sample mode to compute the donor-averaged expression mean and variability for each gene in the transcriptome that had greater than 0.07 mean UMI count. We then performed gene set enrichment analysis using EnrichR to get the significantly enriched gene sets.

### Study subjects and genotyping for Perturb-seq

Our samples were enrolled in PhenoGenetic study (age 18 to 56, average 29.9), as part of the Immvar cohort(20), which were recruited in the Greater Boston Area. Each donor gave written consent to participate and were healthy, without any history of inflammatory disease, autoimmune disease, chronic metabolic disorders or chronic infectious disorders. We genotyped 56 caucasian samples on the OmniExpressExome54 chip, and excluded 2080 SNPs with a call rate < 90% (0.22% of total), 1521 SNPs with Hardy Weinberg *P* < 0.0001 (0.16%) and 259,860 SNPs with MAF < 0.01 (27.04%) out of the total 960,919 SNPs profiled. The Michigan Imputation Server was used to impute these genotypes with the Haplotype Reference Consortium Panel Version r1.1. After genotype imputation had 5,324,560 SNPs, which were then subsetted for our nine donors.

### Regulator target identification and CROP-seq library generation

Our library contained targeted 140 regulators (transcription factors and RNA-binding proteins) with 2 sgRNAs each. Each regulator was unbiasedly chosen using gene expression and accessibility data from activated CD4+ T cells in 95 and 105 healthy donors(18). To get the highly expressed regulators using RNA-seq data, we performed a TMM normalization and took the upper quartile of highly expressed genes and subsetted those that were regulators. To get the regulators with highly accessible binding sites using ATAC-seq data, we enriched for all binding sites on the HOMER database(71) in activated accessible chromatin regions. We took the union of the highly expressed regulators and accessible binding sites, for a total of 140 regulators (Fig. 1B).

The backbone plasmid used to clone the CROP-Seq library was CROPseq-Guide-Puro(28), purchased from Addgene (Addgene. Plasmid #86708). We used two sgRNAs oligo sequences from the Brunello library(88) for each of our chosen 140 regulators. Oligos for the sgRNA library were purchased from Integrated DNA Technologies (IDT) and cloned into the CROPseq plasmid backbone using the methods described by Datlinger et al. [13]. Lentivirus was produced using the UCSF ViraCore.

### SLICE experiment and sequencing

Primary human CD4+ T cells were isolated from peripheral blood mononuclear cells (PBMCs) by magnetic negative selection using the EasySep Human CD4+ T Cell Isolation Kit (STEMCELL, Cat #17952). Cells were cultured in X-Vivo media, consisting of X-Vivo15 medium (Lonza, Cat #04-418Q) with 5% Fetal Calf Serum, 50mM 2-mercaptoethanol, and 10mM N-Acetyl L-Cysteine. On the day of isolation (Day 1), cells were rested in media without stimulation for 24 hours. The day after isolation (Day 2), cells were stimulated with ImmunoCult Human CD3/CD28 T Cell Activator (STEMCELL, Cat #10971) and IL-2 at 50U/mL. 24 hours post stimulation (Day 3), 1 uL of lentivirus was added directly to cultured T cells and gently mixed. Following 24 hours (Day 4), cells were collected, pelleted, and washed in PBS twice. Then, cells were resuspended in Lonza electroporation buffer P3 (Lonza, Cat #V4XP-3032). Cas9 protein (MacroLab, Berkeley, 40mM stock) was added to the cell suspension at a 1:10 v/v ratio. Cells were transferred to a 96 well electroporation cuvette plate (Lonza, cat #VVPA-1002) for nucleofection using the Lonza Nucleofector 96-well Shuttle System and pulse code EH115 (Lonza, cat #VVPA-1002). Immediately after electroporation, pre-warmed media was added to each electroporation well, and 96-well plate was placed at 37 degrees for 20 minutes. Cells were then transferred to culture vessels in X-Vivo media containing 50U/mL IL-2 at 1e6 cells /mL in appropriate tissue culture vessels. Two days later, 1.5ug/mL Puromycin was added in culture media for selection. Cells were expanded every two days, adding fresh media with IL-2 at 50U/mL. Cells were maintained at a cell density of 1e6 cells /mL. On the final day (Day 13) of the experiment, cells from each of the nine donors were counted using Vi-CELL XR and pooled at equal numbers to obtain a final 180,000 cells in 60 uL of PBS. The pooled cells were then processed by UCSF Institute for Human Genetics (IHG) Genomics Core using 16 wells of 10X Chromium Single Cell v2 (PN-120237), as per manufacturer’s protocol, with each well being separately index. The final library was sequenced on two lanes on the Nova-seq for a total of 6.7B reads. To maximize the probability of detecting sgRNAs in cells, we further amplified and sequenced the sgRNA transcripts %from the 10X cDNA library to near saturation as previously described [61] (98%).

### Visualizing gene regulatory networks

To generate the GRNs in 5E, we first used a list of pairs of regulator to their differential-mean expressed genes to define a bipartite graph, which was then visualized in Cytoscape. We then added the connections between the interacting pairs of regulators discovered by differentially correlated genes (DCGs) in the same previously visualized network (5F).

### Identifying candidate interactions for differential correlation analysis

For a transcriptional regulator TR, we first identified all of the DMGs where the TR acts as a transcriptional activator, with the DM coefficient less than 0 across the KO. We then computed the correlation between each TR-DMG pair in WT cells, and constructed the final set of TR-DMG pairs by selected those that had a significant correlation in WT (*ρ* > 0.1). For each of these TR-DMG pairs, we tested for differential correlation across various sgRNAs targeting transcriptional regulators other than TR. The final set of interactions were called by filtering for FDR < 0.1.

### Counting genes with shared TFBS for pairs of transcription factors

For a pair of transcriptional factors TF1 and TF2, we first identified their transcription binding sites (TFBSs) during the ChIP-seq data in the ENCODE datasets. We then took the locations of known gene transcriptional start sites (TSSs) and measured the distance of the nearest TFBS for each TF for each TSS. We then counted the number of genes that have TFBSs of both TF1 and TF2 within a series of window sizes near the TSS, ranging from 10 base pairs to 100K basepairs. We performed this procedure for pairs of TFs chosen at random and also pairs of TFs identified as interacting using differential correlation analysis.

### Assessing the tonic sensitivity of ISGs

We used tonic sensitivty measurements from Gough et al. where the authors compared the expression of ISGs in IFNAR1-KO and WT macrophages [45]. The fold-change between those two groups were defined as the tonic sensitivity, which is the number we use in Fig. 4D.

### eQTL discovery using pseudobulk approach and memento

We used the single cell dataset generated by Perez et al. that profiled peripheral blood mononuclear cells in individuals with systemic lupus erythematosis (SLE) and healthy controls. We maintained the same cell type classifications used in that study.

To identify eQTLs using the psedubulk approach, we first created pseuobulks at the cell-type and individual level by normalizing each cell expression with total UMI count per cell, taking the average for each gene across all individuals, and computing *log*(*x* + 1) for each mean. We filtered genes that had a lower than 0.01 mean UMI counts in the single cell dataset.

We ran Matrix eQTL for each of the Asian and European populations separately, using the same set of genotypes and covariates used by Perez et al. [17]. For `memento`, we also performed the test separately for the two populations, using the same genotypes and covariates. We used the hierarchial resampling mode for `memento`.

### Enrichment of eQTLs in ATAC peaks

We used the same set of ATAC peaks used by Perez et al. [17]. For each SNP, we labeled whether that a cell type specific ATAC peak covered the location of the SNP. We then compared the p-values of the eQTL candidates in a cell-type peak to those of the candidates outside of ATAC peaks using the Wilcoxon Rank Sum test.

### Comparison of eQTLs with OneK1K cohort

To compute the ROC curve and perform power analysis in 6, we compared the eQTLs we discovered using the two approaches to the eQTLs reported by Yazar et al. [18]. We used this much larger dataset as the gold standard to compare methodologies applied to the SLE dataset.

## References

[1] H H McAdams and A Arkin. “Stochastic mechanisms in gene expression”. In: Proc. Natl. Acad. Sci. U. S. A. 94.3 (1997), pp. 814–819. ISSN: 0027-8424. doi: 10.1073/pnas.94.3.814.

[2] Arjun Raj and Alexander van Oudenaarden. “Nature, nurture, or chance: stochastic gene expression and its consequences”. In: Cell 135.2 (2008), pp. 216–226. ISSN: 0092-8674. doi: 10.1016/j.cell.2008.09.050.

[3] Jitao Guo and Xuyu Zhou. Regulatory T cells turn pathogenic. 2015. doi: 10.1038/cmi.2015.12.

[4] John R S Newman et al. “Single-cell proteomic analysis of S. cerevisiae reveals the architecture of biological noise”. In: Nature 441.7095 (2006), pp. 840–846. ISSN: 0028-0836. doi: 10.1038/nature04785.

[5] Arjun Raj et al. “Variability in gene expression underlies incomplete penetrance”. In: Nature 463.7283 (2010), pp. 913–918. ISSN: 0028-0836. doi: 10.1038/nature08781.

[6] Avigdor Eldar and Michael B. Elowitz. Functional roles for noise in genetic circuits. 2010. doi: 10.1038/nature09326.

[7] Maike M K Hansen et al. “Cytoplasmic Amplification of Transcriptional Noise Generates Substantial Cell-to-Cell Variability”. In: Cell Syst 7.4 (2018), 384–397.e6. ISSN: 2405-4712. doi: 10.1016/j.cels.2018.08.002.

[8] Ido Golding et al. “Real-time kinetics of gene activity in individual bacteria”. In: Cell 123.6 (2005), pp. 1025–1036. ISSN: 0092-8674. doi: 10.1016/j.cell.2005.09.031.

[9] Brian Munsky, Gregor Neuert, and Alexander van Oudenaarden. “Using gene expression noise to understand gene regulation”. In: Science 336.6078 (2012), pp. 183–187. ISSN: 0036-8075. doi: 10.1126/science.1216379.

[10] Anika Gupta et al. “Inferring gene regulation from stochastic transcriptional variation across single cells at steady state”. en. In: Proc. Natl. Acad. Sci. U. S. A. 119.34 (Aug. 2022), e2207392119.

[11] Ker-Chau Li. “Genome-wide coexpression dynamics: theory and application”. en. In: Proc. Natl. Acad. Sci. U. S. A. 99.26 (Dec. 2002), pp. 16875–16880.

[12] Sanjay R. Srivatsan et al. “Massively multiplex chemical transcriptomics at single-cell resolution”. In: Science 367.6473 (2020), pp. 45–51. ISSN: 10959203. doi: 10.1126/science.aax6234.

[13] Paul Datlinger et al. “Ultra-high throughput single-cell RNA sequencing by combinatorial fluidic indexing”. In: bioRxiv (2019), pp. 1–27. doi: 10.1101/2019.12.17.879304.

[14] Atray Dixit et al. “Perturb-Seq: Dissecting Molecular Circuits with Scalable Single-Cell RNA Profiling of Pooled Genetic Screens”. In: Cell 167.7 (2016), 1853–1866.e17. ISSN: 10974172. doi: 10.1016/j.cell.2016.11.038.

[15] Hyun Min Kang et al. “Multiplexed droplet single-cell RNA-sequencing using natural genetic variation”. In: Nature Biotechnology 36.1 (2018), pp. 89–94. ISSN: 15461696. doi: 10.1038/nbt.4042.

[16] Monique G.P. Van Der Wijst et al. “Single-cell RNA sequencing identifies celltype-specific cis-eQTLs and co-expression QTLs”. In: Nature Genetics 50.4 (2018), pp. 493–497. ISSN: 15461718. doi: 10.1038/s41588-018-0089-9.

[17] Richard K Perez et al. “Single-cell RNA-seq reveals cell type-specific molecular and genetic associations to lupus”. In: Science 376.6589 (2022), eabf1970. ISSN: 0036-8075. doi: 10.1126/science.abf1970.

[18] Seyhan Yazar et al. “Single-cell eQTL mapping identifies cell type-specific genetic control of autoimmune disease”. en. In: Science 376.6589 (2022), eabf3041.

[19] Jordan W Squair et al. “Confronting false discoveries in single-cell differential expression”. In: Nat. Commun. 12.1 (2021), p. 5692. ISSN: 2041-1723. doi: 10.1038/s41467-021-25960-2.

[20] Charlotte Soneson and Mark D Robinson. “Bias, robustness and scalability in single-cell differential expression analysis”. In: Nat. Methods 15.4 (2018), pp. 255–261. ISSN: 1548-7091. doi: 10.1038/nmeth.4612.

[21] David Lähnemann et al. “Eleven grand challenges in single-cell data science”. In: Genome Biol. 21.1 (2020), p. 31. ISSN: 1465-6906. doi: 10.1186/s13059-020-1926-6.

[22] Abhishek Sarkar and Matthew Stephens. “Separating measurement and expression models clarifies confusion in single-cell RNA sequencing analysis”. In: Nat. Genet. 53.6 (2021), pp. 770–777. ISSN: 1061-4036. doi: 10.1038/s41588-021-00873-4.

[23] Romain Lopez et al. “Deep generative modeling for single-cell transcriptomics”. In: Nature Methods 15.12 (2018), pp. 1053–1058. ISSN: 15487105. doi: 10.1038/s41592-018-0229-2.

[24] Davide Risso et al. “A general and flexible method for signal extraction from single-cell RNA-seq data”. In: Nature Communications 9.1 (2018), pp. 1–17. ISSN: 20411723. doi: 10.1038/s41467-017-02554-5.

[25] Keegan D Korthauer et al. “A statistical approach for identifying differential distributions in single-cell RNA-seq experiments”. In: Genome Biol. 17.1 (2016), p. 222. ISSN: 1465-6906. doi: 10.1186/s13059-016-1077-y.

[26] Greg Finak et al. “MAST: A flexible statistical framework for assessing transcriptional changes and characterizing heterogeneity in single-cell RNA sequencing data”. In: Genome Biology 16.1 (2015), p. 278. ISSN: 1474760X. doi: 10.1186/s13059-015-0844-5.

[27] Nils Eling et al. “Correcting the Mean-Variance Dependency for Differential Variability Testing Using Single-Cell RNA Sequencing Data.” In: Cell systems 7.3 (2018), 284–294.e12. ISSN: 2405-4712. doi: 10.1016/j.cels.2018.06.011.

[28] Christopher S McGinnis et al. “MULTI-seq: sample multiplexing for single-cell RNA sequencing using lipid-tagged indices”. In: Nat. Methods 16.7 (2019), pp. 619–626. ISSN: 1548-7091. doi: 10.1038/s41592-019-0433-8.

[29] Marlon Stoeckius et al. “Cell Hashing with barcoded antibodies enables multiplexing and doublet detection for single cell genomics”. In: Genome Biology 19.1 (2018), p. 224. ISSN: 1474760X. doi: 10.1186/s13059-018-1603-1.

[30] Junyue Cao et al. “Comprehensive single-cell transcriptional profiling of a multicellular organism”. In: Science 357.6352 (2017), pp. 661–667. ISSN: 0036-8075. doi: 10.1126/science.aam8940.

[31] F. Alexander Wolf, Philipp Angerer, and Fabian J. Theis. “SCANPY: Large-scale single-cell gene expression data analysis”. In: Genome Biology 19.1 (2018), p. 15. ISSN: 1474760X. doi: 10.1186/s13059-017-1382-0.

[32] Yu Fu et al. “Elimination of PCR duplicates in RNA-seq and small RNA-seq using unique molecular identifiers”. In: BMC Genomics 19.1 (2018), p. 531. ISSN: 1471-2164. doi: 10.1186/s12864-018-4933-1.

[33] Allon M. Klein et al. “Droplet barcoding for single-cell transcriptomics applied to embryonic stem cells”. In: Cell 161.5 (2015), pp. 1187–1201. ISSN: 10974172. doi: 10.1016/j.cell.2015.04.044.

[34] Dominic Grün, Lennart Kester, and Alexander Van Oudenaarden. “Validation of noise models for single-cell transcriptomics”. In: Nat. Methods 11.6 (2014), pp. 637–640. ISSN: 1548-7091. doi: 10.1038/nmeth.2930.

[35] Shila Ghazanfar et al. “Investigating higher-order interactions in single-cell data with scHOT”. In: Nat. Methods 17.8 (2020), pp. 799–806. ISSN: 1548-7091. doi: 10.1038/s41592-020-0885-x.

[36] Martin Jinye Zhang, Vasilis Ntranos, and David Tse. “Determining sequencing depth in a single-cell RNA-seq experiment”. In: Nat. Commun. 11.1 (2020), pp. 1–11. ISSN: 2041-1723. doi: 10.1038/s41467-020-14482-y.

[37] Michael Hagemann-Jensen et al. “Single-cell RNA counting at allele and isoform resolution using Smart-seq3”. In: Nature Biotechnology 38.6 (2020), pp. 708–714. ISSN: 15461696. doi: 10.1038/s41587-020-0497-0.

[38] Eduardo Torre et al. “Rare Cell Detection by Single-Cell RNA Sequencing as Guided by Single-Molecule RNA FISH”. In: Cell Syst 6.2 (2018), 171–179.e5. ISSN: 2405-4712. doi: 10.1016/j.cels.2018.01.014.

[39] Mo Huang et al. “SAVER: Gene expression recovery for single-cell RNA sequencing”. In: Nat. Methods 15.7 (2018), pp. 539–542. ISSN: 1548-7091. doi: 10.1038/s41592-018-0033-z.

[40] Bradley. Efron and Robert. Tibshirani. An introduction to the bootstrap. Chapman & Hall, 1994, p. 436. ISBN: 9780412042317.

[41] Rishi R Goel, Sergei V Kotenko, and Mariana J Kaplan. “Interferon lambda in inflammation and autoimmune rheumatic diseases”. en. In: Nat. Rev. Rheumatol. 17.6 (June 2021), pp. 349–362.

[42] Liqun Zhang et al. “Infection of ciliated cells by human parainfluenza virus type 3 in an in vitro model of human airway epithelium”. In: J. Virol. 79.2 (2005), pp. 1113–1124. ISSN: 0022-538X. doi: 10.1128/JVI.79.2.1113-1124.2005.

[43] Nai-Huei Wu et al. “The differentiated airway epithelium infected by influenza viruses maintains the barrier function despite a dramatic loss of ciliated cells”. In: Sci. Rep. 6 (2016), p. 39668. ISSN: 2045-2322. doi: 10.1038/srep39668.

[44] Neal G Ravindra et al. “Single-cell longitudinal analysis of SARS-CoV-2 infection in human airway epithelium identifies target cells, alterations in gene expression, and cell state changes”. In: PLoS Biol. 19.3 (2021), e3001143. ISSN: 1544-9173. doi: 10.1371/journal.pbio.3001143.

[45] Daniel J. Gough et al. Constitutive Type I Interferon Modulates Homeostatic Balance through Tonic Signaling. 2012. doi: 10.1016/j.immuni.2012.01.011.

[46] Konrad C Bradley et al. “Microbiota-Driven Tonic Interferon Signals in Lung Stromal Cells Protect from Influenza Virus Infection”. In: Cell Rep. 28.1 (2019), 245–256.e4. ISSN: 2211-1247. doi: 10.1016/j.celrep.2019.05.105.

[47] Tzachi Hagai et al. “Gene expression variability across cells and species shapes innate immunity”. en. In: Nature 563.7730 (Nov. 2018), pp. 197–202.

[48] Sara Mostafavi et al. “Parsing the Interferon Transcriptional Network and Its Disease Associations In Brief Resource Parsing the Interferon Transcriptional Network and Its Disease Associations”. In: (2016). doi: 10.1016/j.cell.2015.12.032.

[49] Rachel E Gate et al. “Genetic determinants of co-accessible chromatin regions in activated T cells across humans”. In: Nat. Genet. 50.8 (2018), pp. 1140–1150. ISSN: 1061-4036. doi: 10.1038/s41588-018-0156-2.

[50] Ian Dunham et al. “An integrated encyclopedia of DNA elements in the human genome”. In: Nature 489.7414 (2012), pp. 57–74. ISSN: 0028-0836. doi: 10.1038/nature11247.

[51] Aparna Nathan et al. “Single-cell eQTL models reveal dynamic T cell state dependence of disease loci”. en. In: Nature 606.7912 (June 2022), pp. 120–128.

[52] Joseph M Replogle et al. “Mapping information-rich genotype-phenotype landscapes with genome-scale Perturb-seq”. en. In: Cell 185.14 (July 2022), 2559–2575.e28.

[53] F. William Townes et al. “Feature selection and dimension reduction for single-cell RNA-Seq based on a multinomial model”. In: Genome Biology 20.1 (2019), p. 295. ISSN: 1474760X. doi: 10.1186/s13059-019-1861-6.

[54] 10X Genomics. Single cell profiling on more samples with no time constraints. https://pages.10xgenomics.com/rs/446-PBO-704/images/10x_Product-Sheet_LIT000159_Fixed-RNA-Profiling_Letter_Digital.pdf. Accessed: 2022-10-8.

[55] 10X Genomics. Visualize gene expression within the tissue context. https://pages.10xgenomics.com/rs/446-PBO-704/images/10x_LIT059_ProductSheet_VisiumSpatialGeneExpression_Letter_digital.pdf. Accessed: 2022-10-8.

[56] Luyi Tian, Fei Chen, and Evan Z Macosko. “The expanding vistas of spatial transcriptomics”. en. In: Nat. Biotechnol. (Oct. 2022).

[57] Grace X Y Zheng et al. “Massively parallel digital transcriptional profiling of single cells”. In: Nat. Commun. 8.1 (2017), pp. 1–12. ISSN: 2041-1723. doi: 10.1038/ncomms14049.

[58] Mark D. Robinson and Alicia Oshlack. “A scaling normalization method for differential expression analysis of RNA-seq data”. In: Genome Biology 11.3 (2010), R25. ISSN: 14747596. doi: 10.1186/gb-2010-11-3-r25.

[59] Michael I Love, Wolfgang Huber, and Simon Anders. “Moderated estimation of fold change and dispersion for RNA-seq data with DESeq2”. In: Genome Biology 15.12 (2014), p. 550. ISSN: 1474-760X. doi: 10.1186/s13059-014-0550-8.

[60] Aaron T.L. Lun, Karsten Bach, and John C. Marioni. “Pooling across cells to normalize single-cell RNA sequencing data with many zero counts”. In: Genome Biology 17.1 (2016), pp. 1–14. ISSN: 1474760X. doi: 10.1186/s13059-016-0947-7.

[61] Andrew J Hill et al. “On the design of CRISPR-based single-cell molecular screens”. In: Nat. Methods 15.4 (2018), pp. 271–274. ISSN: 1548-7091. doi: 10.1038/nmeth.4604.

