## Supplementary figures for "memento: Generalized differential expression analysis of single-cell RNA-seq with method of moments estimation and efficient resampling"

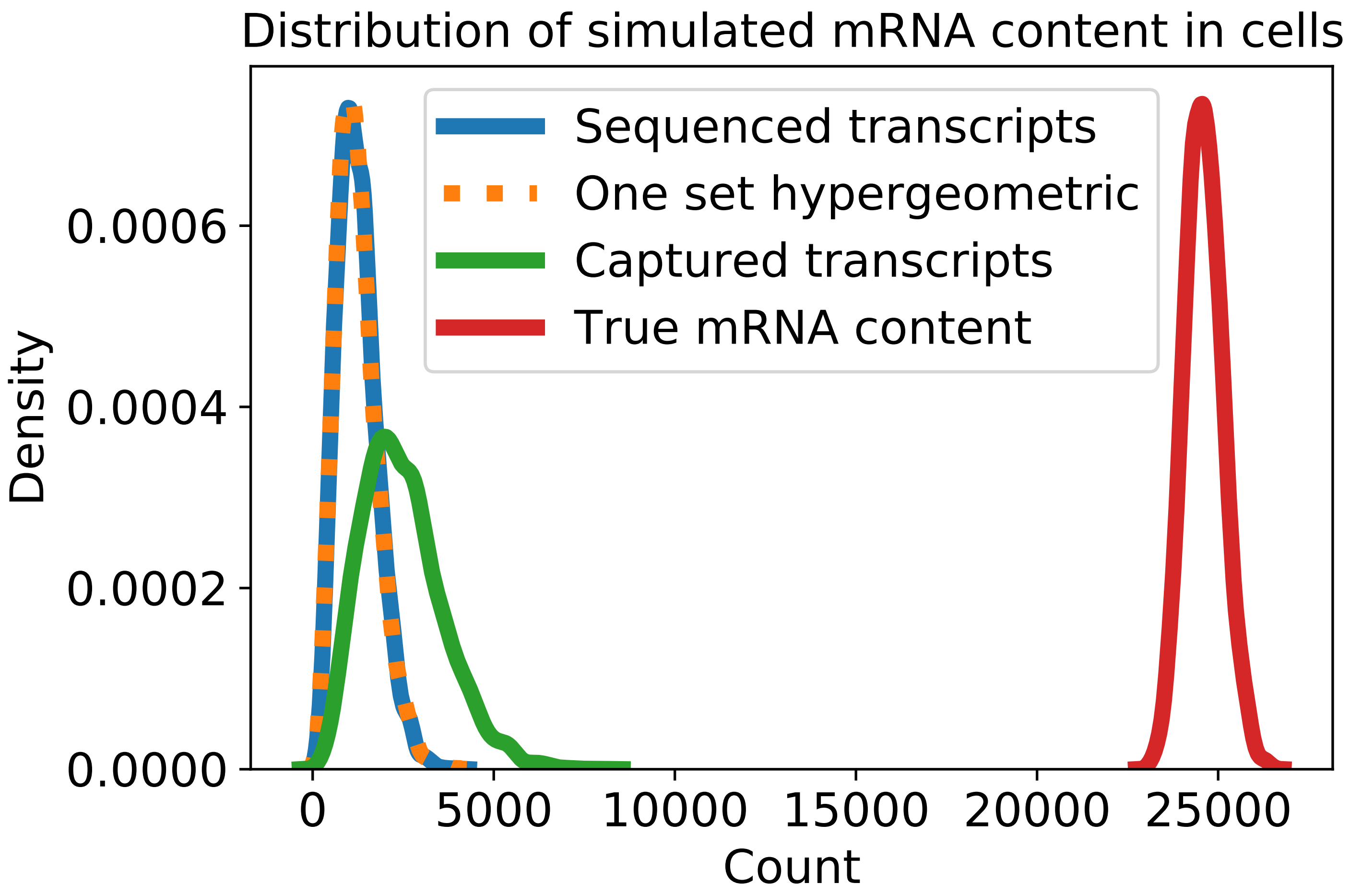


**Figure S1:** Single step of hypergeometric sampling well approximates the compound sampling process from capture and sequencing.

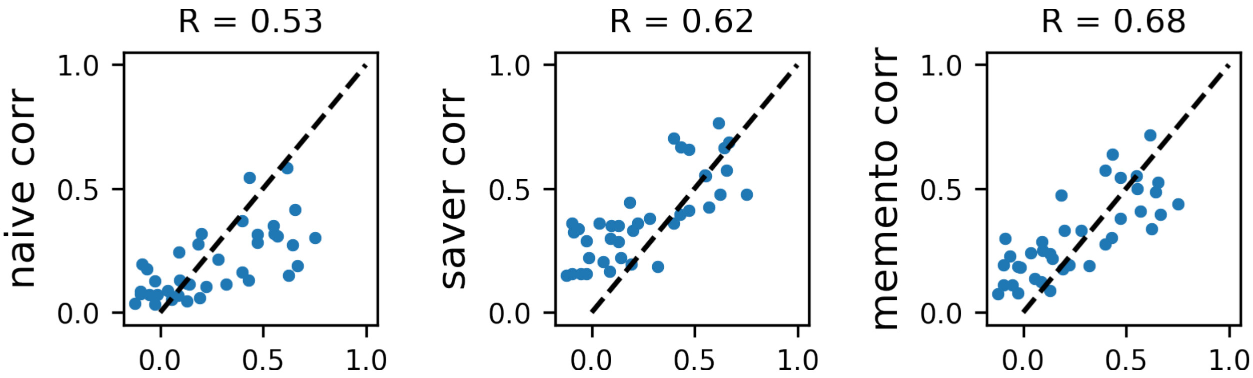


**Figure S2:** Correlations computed using naive, `memento`, and SAVER imputed data (y-axis) against gene correlations from FISH (x-axis)

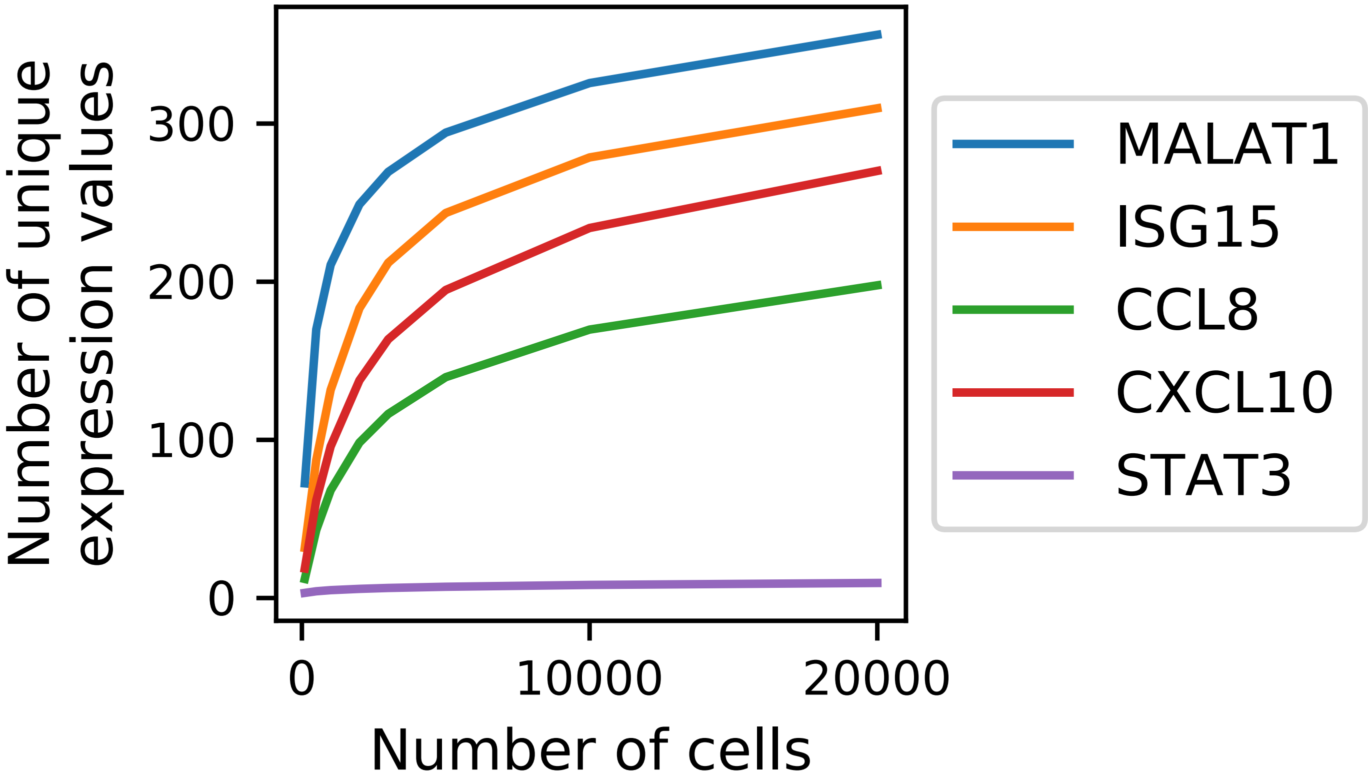


**Figure S3:** Number of unique transcript counts (y-axis) against the number of cells present in the dataset (x-axis)

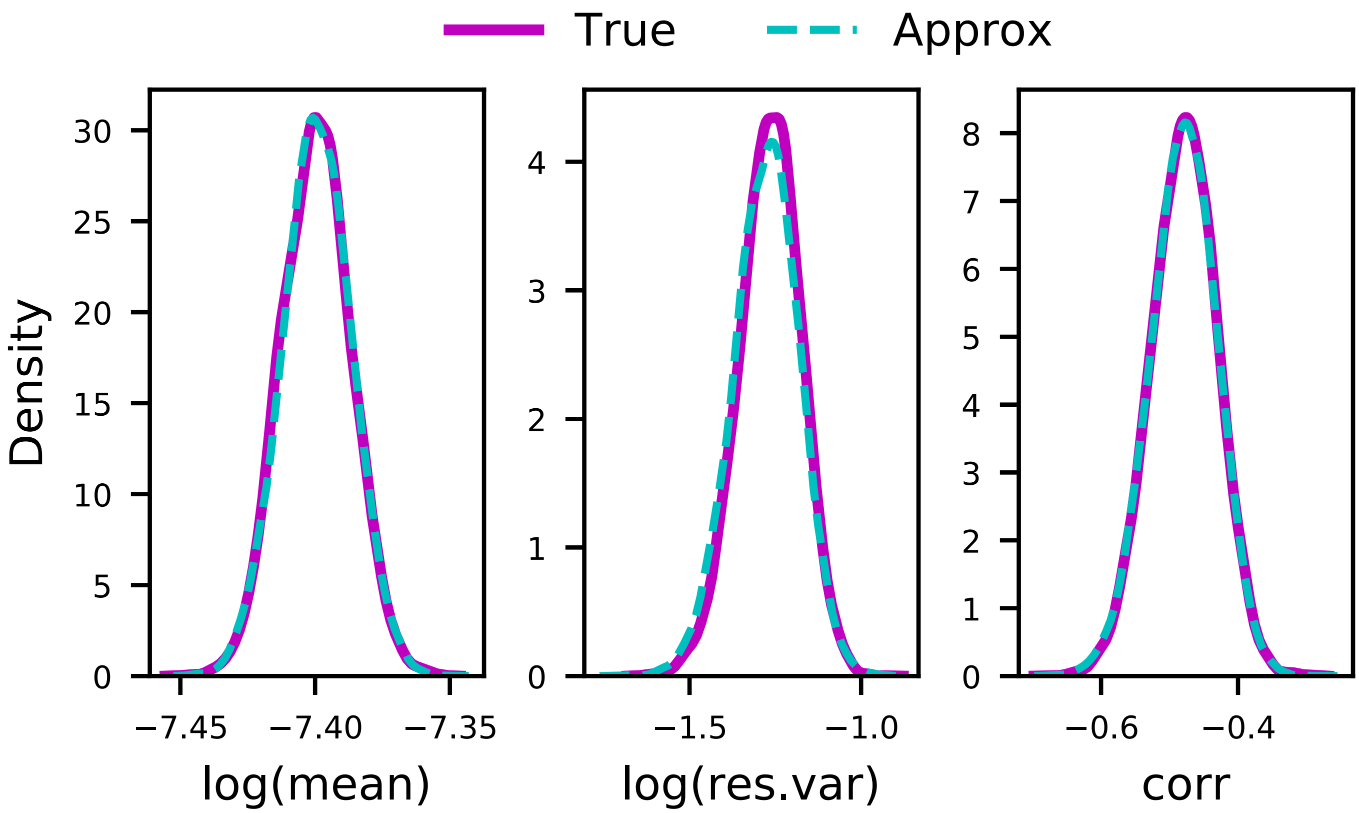


**Figure S4:** Efficient bootstrap in `memento` vs full bootstrap.

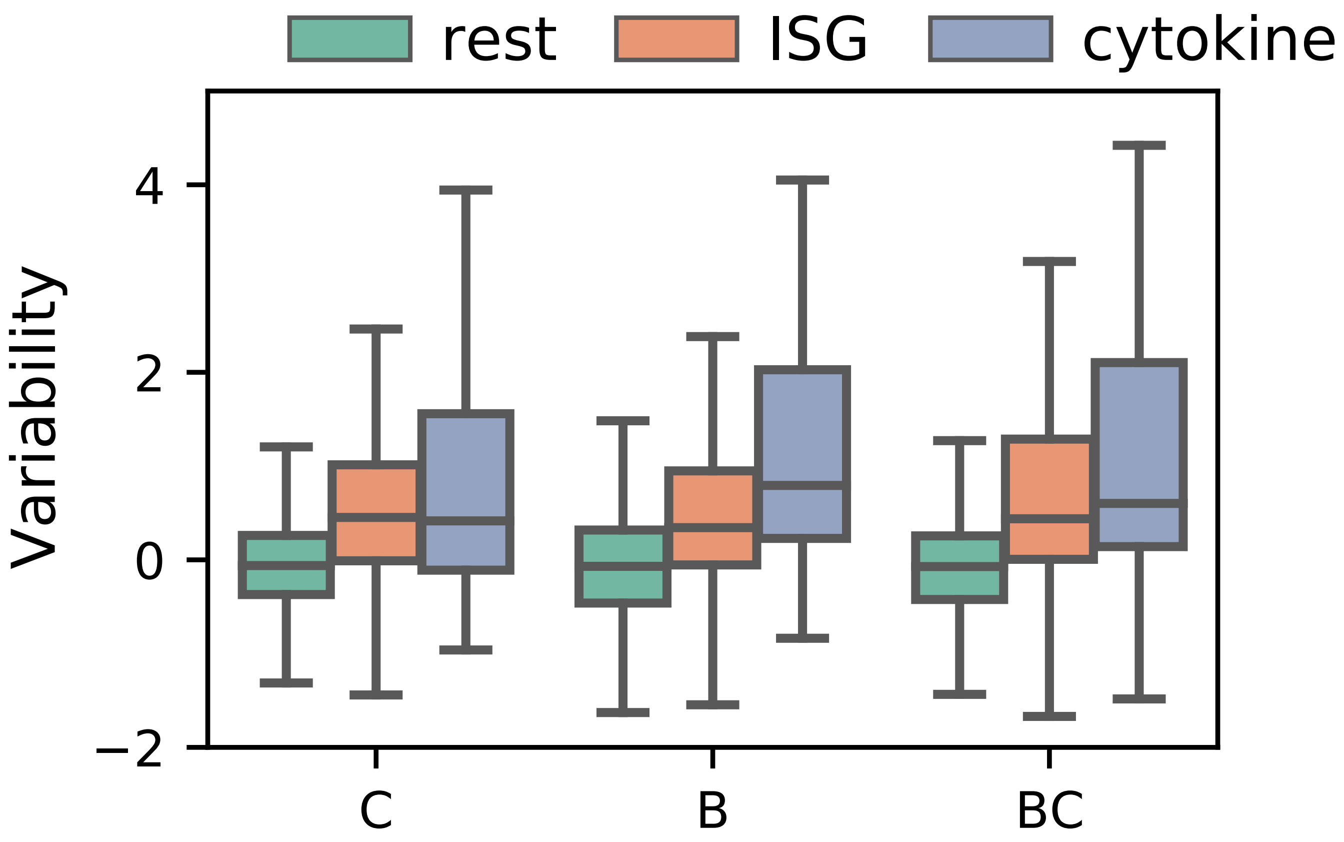


**Figure S5:** Gene expression variability (y-axis) for each class of genes across cell types.

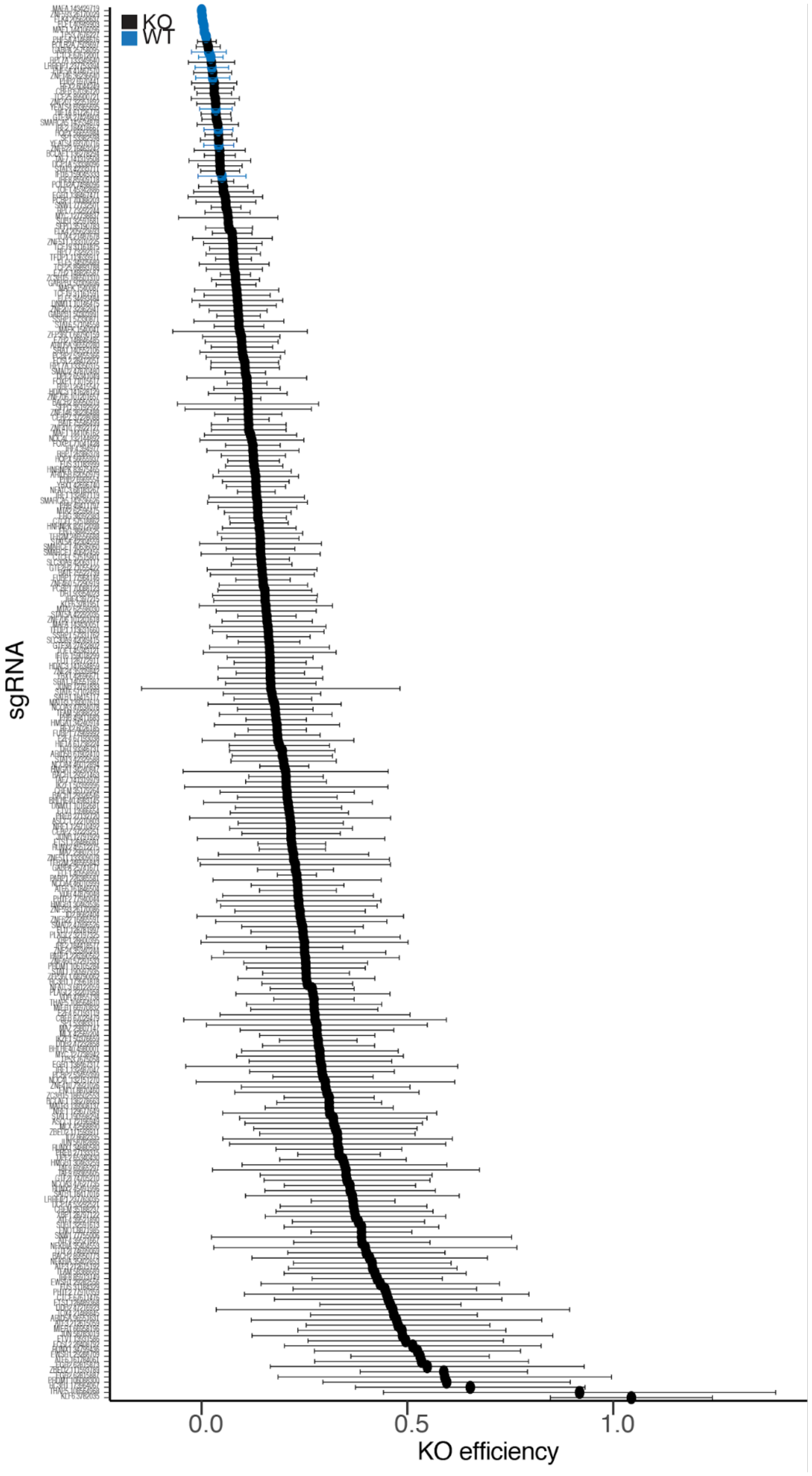


**Figure S6:** We estimated the average sgRNA KO efficiency (x-axis) per sgRNA (y-axis). Each point represents the average KO efficiency and error bars are the standard deviations across donors.

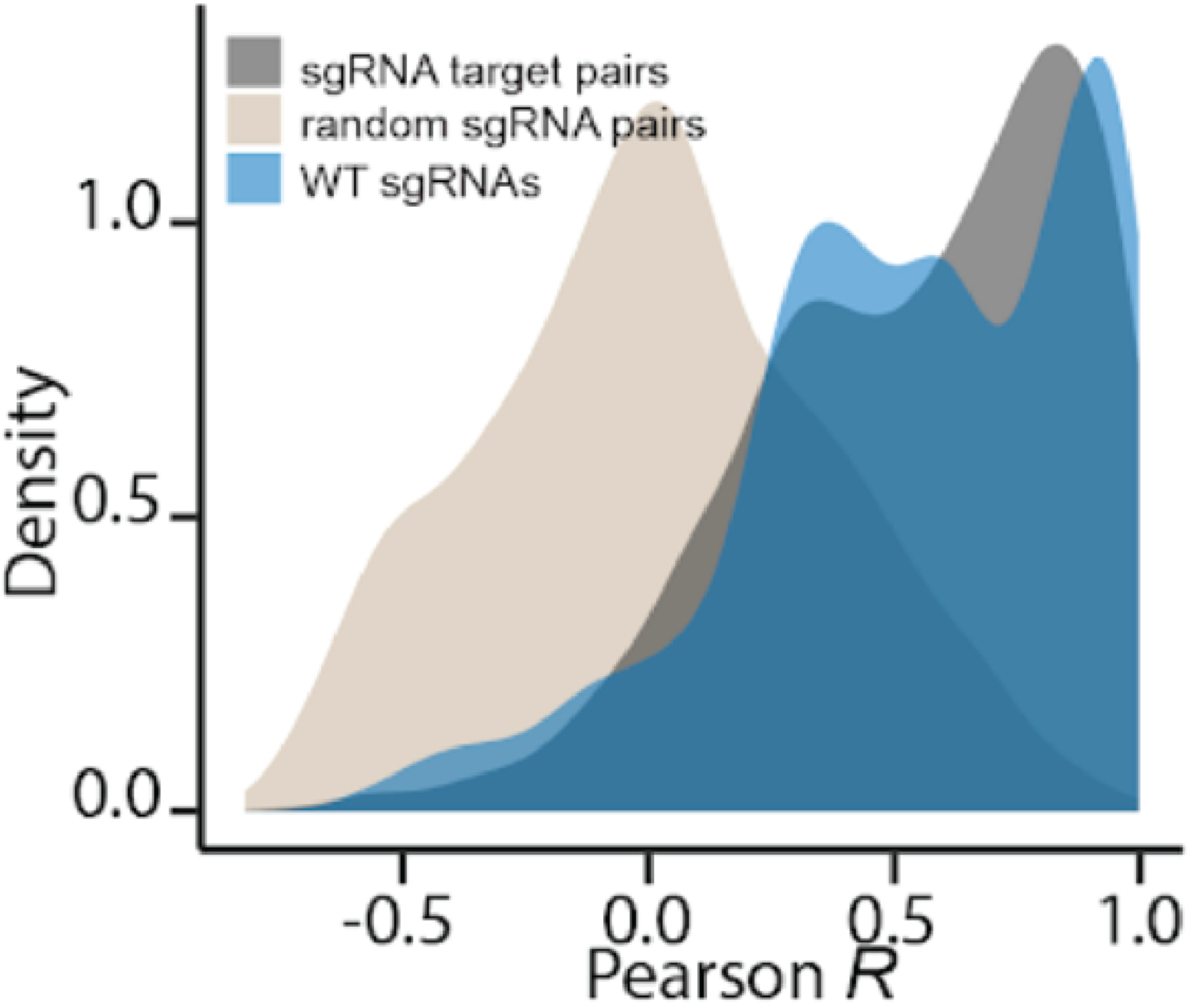


**Figure S7:** Distribution of transcriptome correlations for WT guides, guides targeting the same gene, and random pairs of guides.

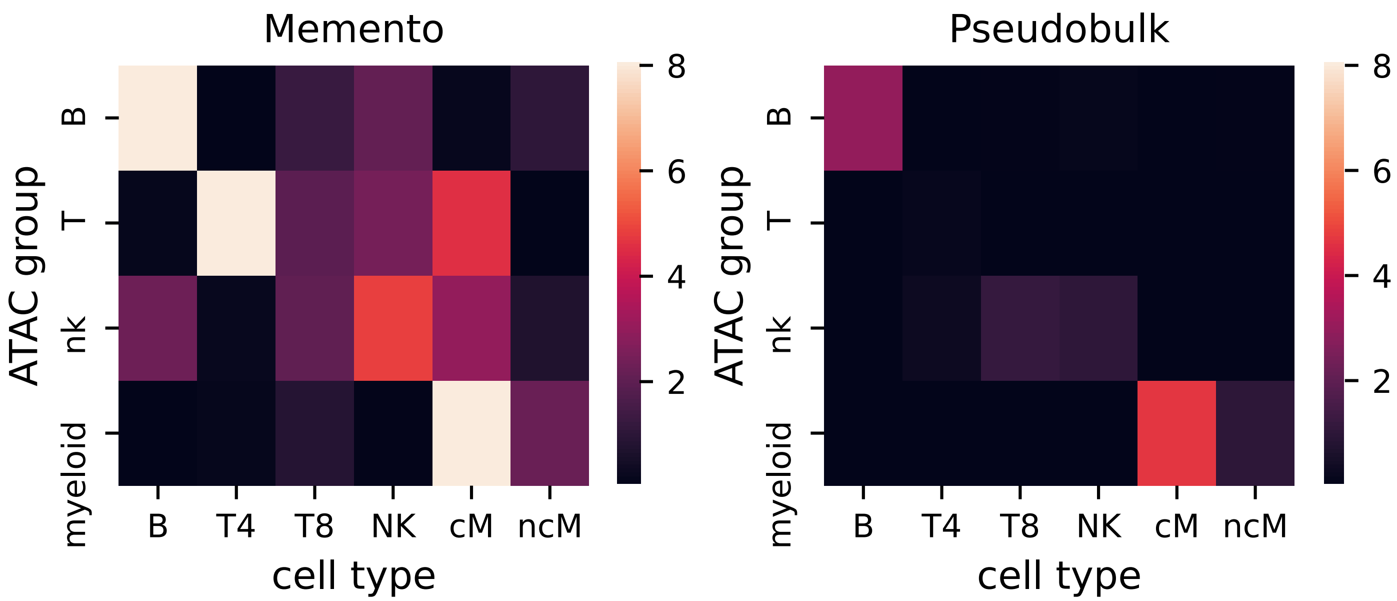


**Figure S8:** Enrichment of eQTLs in cell-type specific ATAC peaks (Europeans).

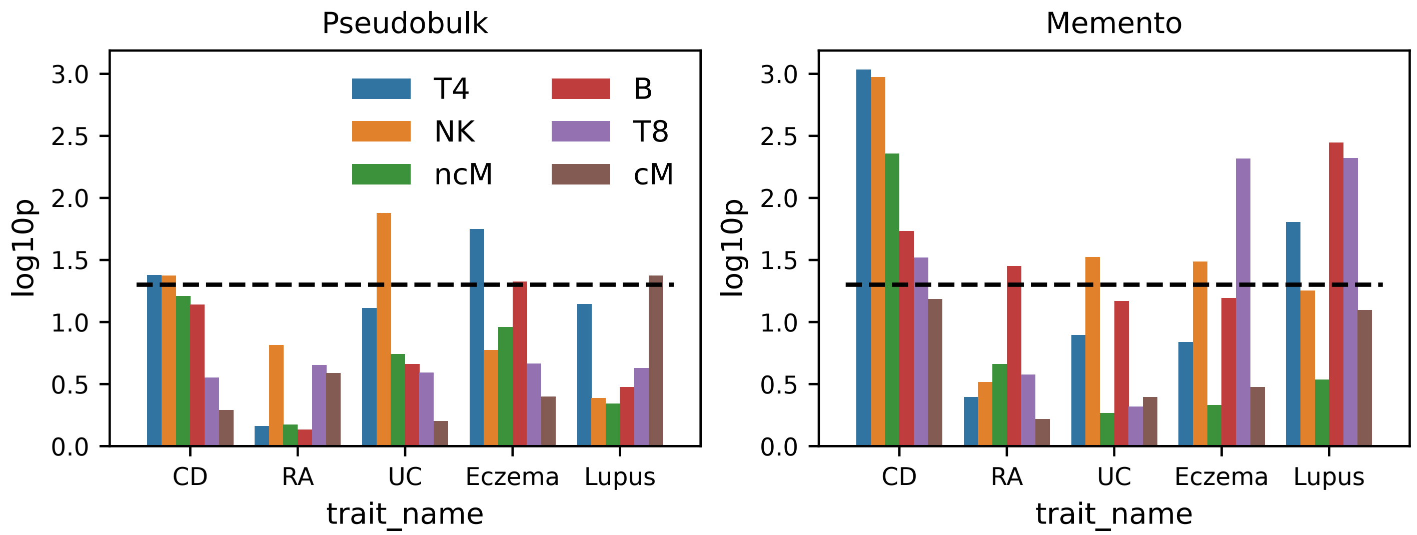


**Figure S9:** LDSC-score regression enrichment for diseases using eGenes found via `memento` and the pseudobulk method.

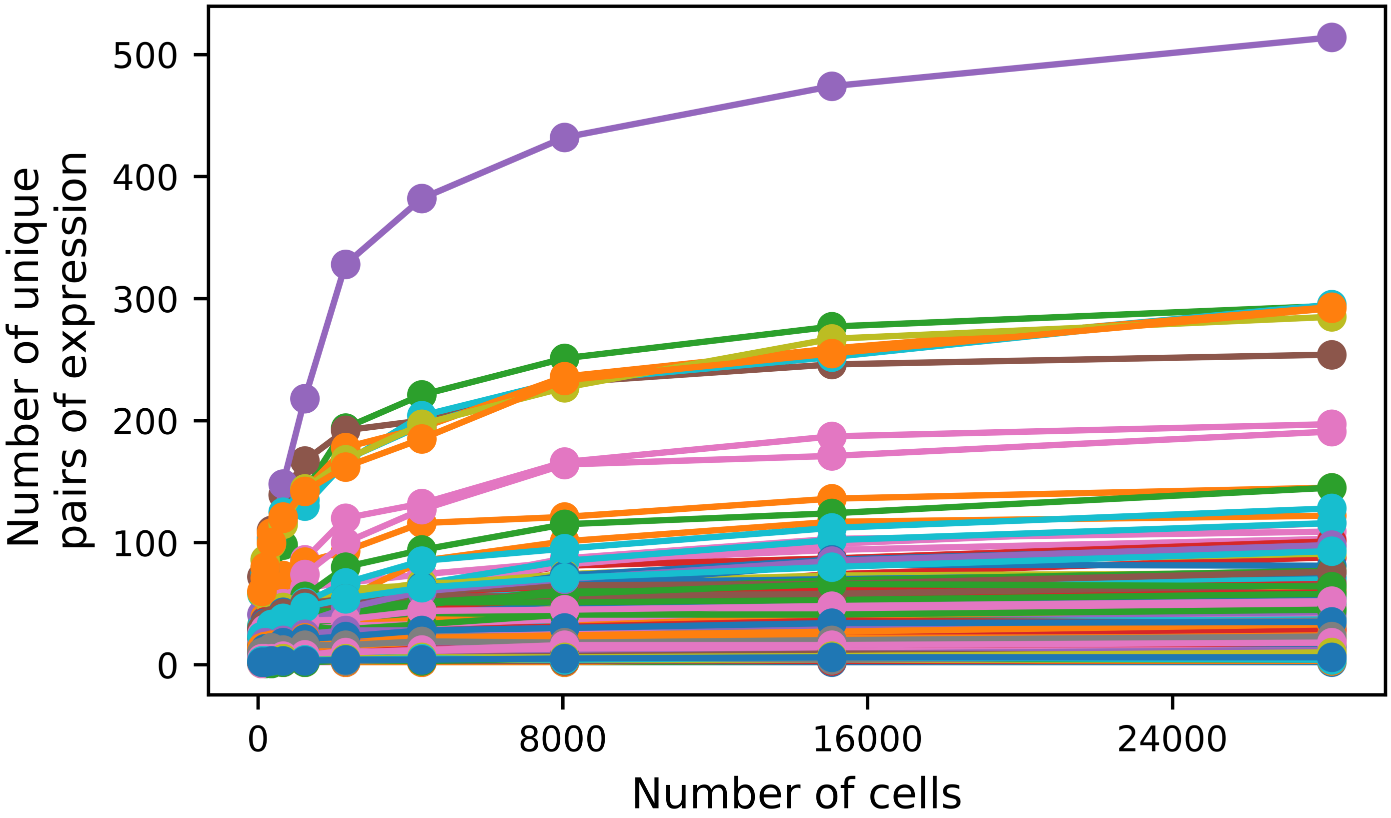


**Figure S10:** Number of unique pairs of genes for randomly selected pairs in the IFN-B dataset.
